# A functional map of phosphoprotein phosphatase regulation identifies an evolutionary conserved reductase for the catalytic metal ions

**DOI:** 10.1101/2025.02.12.637884

**Authors:** Bob Meeusen, Sara M. Ambjørn, Jiri Veis, Rachel C. Riley, Gianmatteo Vit, Brooke L. Brauer, Mads H. Møller, Elora C. Greiner, Camilla B. Chan, Melanie B. Weisser, Dimitriya H. Garvanska, Hao Zhu, Norman E. Davey, Arminja N. Kettenbach, Egon Ogris, Jakob Nilsson

## Abstract

Serine/Threonine phosphoprotein phosphatases (PPPs, PP1-PP7) are conserved metalloenzymes and central to intracellular signaling in eukaryotes, but the details of their regulation is poorly understood. To address this, we performed genome-wide CRISPR knockout and focused base editor screens in PPP perturbed conditions to establish a high-resolution functional map of PPP regulation that pinpoints novel regulatory mechanisms. Through this, we identify the orphan reductase CYB5R4 as an evolutionarily conserved activator of PP4 and PP6, but not the closely related PP2A. Heme binding is essential for CYB5R4 function and mechanistically involves the reduction of the metal ions in the active site. Importantly, CYB5R4-mediated activation of PP4 is critical for cell viability when cells are treated with DNA damage-inducing agents known to cause oxidative stress. The discovery of a dedicated PPP reductase points to shared regulatory principles with protein tyrosine phosphatases, where specific enzymes dictate activity by regulating the active site redox state. In sum, our work provides a resource for understanding PPP function and the regulation of intracellular signaling.

## Introduction

Reversible protein phosphorylation is an essential regulatory mechanism of cells to respond to extra- and intracellular cues. The interplay between protein kinases and protein phosphatases controls the phosphorylation state of tyrosine, serine and threonine residues on their target proteins, collectively dictating the outputs from signaling pathways (1). Phosphoprotein phosphatases (PPPs, PP1-PP7) constitute a highly conserved family of serine/threonine protein phosphatases, accounting for the majority of dephosphorylation activity in eukaryotic cells (2). The PPP catalytic subunits incorporate two metal ions to activate a water molecule that acts as the nucleophile in the dephosphorylation reaction (3). Despite a highly similar catalytic core structure, the PPP family members achieve specificity by incorporating their catalytic subunits into family-specific holoenzymes. The PP2A, PP4 and PP6 catalytic subunits (referred to as PP2A C, PP4 C, PP6 C) uniquely form stable trimeric holoenzymes with a structural subunit and a substrate-specifying regulatory subunit (4,5). These PP2A-like phosphatases are widely involved in core biological processes, such as mitogenic signaling, cell cycle regulation and the DNA damage response, and consistent with this, dysregulation of these proteins is linked to human diseases (6–9). Central to the regulation of the PP2A-like phosphatase holoenzymes is the tightly orchestrated assembly and activation, which involves the action of at least five regulatory proteins (alpha4, PTPA, LCMT1, PME-1 and TIPRL), governing catalytic metal ion coordination and trimeric holoenzyme composition (**Fig. 1A**) (4,5,10–15). However, many aspects of PP2A-like phosphatase assembly and activation are poorly understood, despite being fundamental for understanding phospho-dependent signaling in eukaryotes.

**Figure 1:**
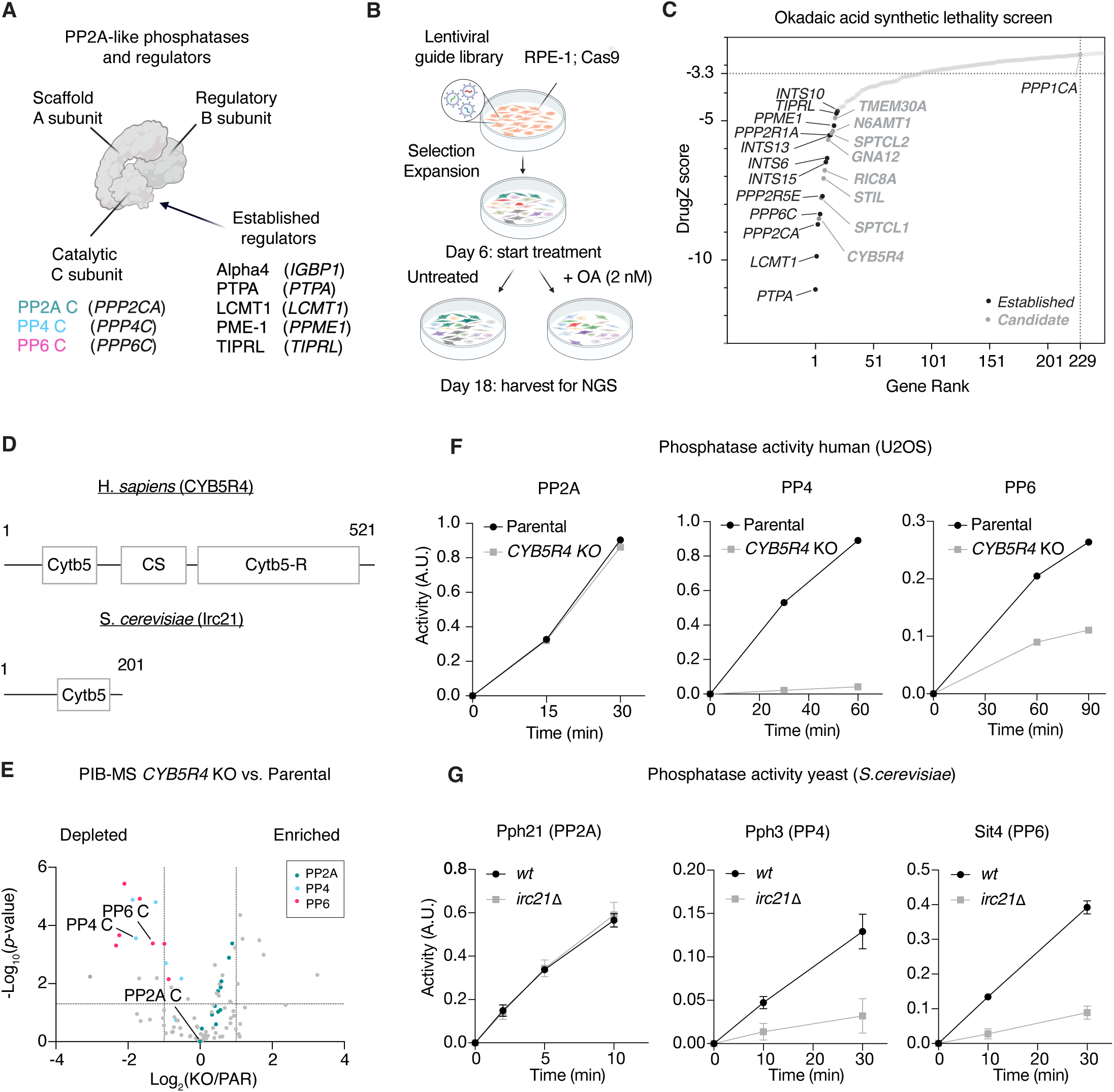
Genome-wide CRISPR-Cas9 knockout screen identifies CYB5R4 as an evolutionarily conserved PPP regulator. A) Holoenzyme composition and regulators of PP2A-like phosphatases. B) Schematic of genome-wide CRISPR-Cas9 screen for genes whose knockout are synthetic lethal with okadaic acid (OA). NGS, next-generation sequencing. C) DrugZ analysis of CRISPR-Cas9 synthetic lethality screen performed in RPE1 p53-/-cells comparing treatment with a low dose of OA (2 nM) and untreated conditions. The top genes are indicated with established PP2A-like holoenzyme components and regulators in black and candidate regulators in grey. Significance threshold (*p*-value of 0.05) is indicated with a dotted line at a DrugZ score of −3.3. *PPP1CA* is indicated as highest scoring non PP2A-like phosphatase. D) Human CYB5R4 and yeast Irc21 domain organization. Cytb5, cytochrome B5 domain. CS, CHORD/SGT domain. Cytb5-R, Cytochrome b5 reductase domain. E) Profiling of PPP composition by phosphatase inhibitor beads and mass spectrometry (PIB-MS). Volcano plot comparing the phosphatase components captured on phosphatase inhibitor beads from U2OS *CYB5R4* knockout (KO) and parental (PAR) cells. PP2A holoenzyme components are indicated in green, PP4 in blue and PP6 in pink. ‘C’ indicates catalytic subunit. F) Peptide dephosphorylation assays measuring the activity of 3xFLAG-tagged PP2A C, PP4 C, and PP6 C, immunopurified from U2OS parental or *CYB5R4* knockout stable cell lines. The data is representative of three independent experiments. G) Peptide dephosphorylation assays measuring the activity of HA-Pph21, myc-Pph3, and HA-Sit4 immunopurified from endogenously tagged wildtype (*wt*) and *irc21* deletion strains. Data is shown for three independent experiments, and error bars represent standard deviations.

## Results

### A genome-wide CRISPR knockout screen identifies CYB5R4 as an evolutionary conserved activator of PP4 and PP6 protein phosphatases

To expand our understanding of PP2A-like phosphatase regulation, we performed a genome-wide CRISPR-Cas9 knockout synthetic lethality screen in RPE1 p53-/-cells to potentially identify additional regulatory mechanisms (**Fig. 1B**) (16). We reasoned that in the presence of a low dose of the PPP inhibitor okadaic acid (OA), which at 2 nM primarily inhibits PP2A-like phosphatases (17,18), the knockout of a gene required for PP2A-like phosphatase activity would be synthetic lethal due to strongly reduced phosphatase activity. Our screening approach was validated by the fact that the top synthetic lethal genes included PP2A-like holoenzyme components (*PPP2CA*, *PPP2R1A*, *PPP2R5E*, *PPP6C, INTS6/10/13/15*), and importantly, four out of five regulators of holoenzyme assembly (*PTPA*, *LCMT1*, *PPME1*, *TIPRL*) (**Fig. 1C, Table S1**). Among the top 20 synthetic lethal genes, we additionally identified eight candidates with no or poorly understood roles in PPP regulation (19–22), which we here refer to as candidate phosphatase regulators. These eight candidates covered several distinct biological functions, including reductase activity (*CYB5R4*) (23–25), ceramide synthesis (*SPTCL1* and *SPTLC2*) (26,27), and extracellular signal transduction (GTPase complex *RIC8A* and *GNA12*) (28,29) (**Fig. 1C**).

To confirm that our screen indeed had identified novel phosphatase regulators, we focused our attention on the top scoring candidate, the cytochrome b5 oxidoreductase CYB5R4. CYB5R4 is a multidomain protein consisting of a Cytochrome b5 (Cytb5) heme-binding domain linked with a CS domain to its Cytochrome b5 reductase domain (Cytb5-R) (**Fig. 1D**), which can reduce various substrates *in vitro* (23–25). Although previously linked to diabetes and oxidative stress in mouse models (30,31) and genetically correlated with the PP2A-like phosphatase regulators PTPA and TIPRL (19,32), its cellular function remains largely unknown.

To validate our screen results, we generated U2OS *CYB5R4* knockout cells and confirmed that they are hypersensitive to OA, and this effect could be rescued by reintroduction of CYB5R4 (**Fig. S1A-C**). Since OA can target multiple PPPs in addition to PP2A, PP4, and PP6 (although at higher IC_50_’s), we next sought to get an unbiased view of which PPPs were affected by loss of CYB5R4. To this end, we used the phosphatase inhibitor beads and mass spectrometry (PIB-MS) approach, where all PPP catalytic subunits and their associated proteins are captured on microcystin-LR beads (MC-beads) followed by quantitative MS analysis, allowing for unbiased PPP profiling (33). Capturing PPPs on MC-beads from lysates of U2OS parental and *CYB5R4* KO cells revealed the specific loss of PP4 and PP6 but not PP2A holoenzyme components in *CYB5R4* KO cells, which was rescued by reintroduction of CYB5R4 (**Fig. 1E, Fig. S1D, Table S2**). The specific loss of PP4 and PP6 components was reproducible in RPE-1 cells, MEFs from *CYB5R4* knockout mice (30), observed at physiological oxygen levels, and not due to changes at the proteome level (**Fig. S1E-G, Table S2-6**). As microcystin-LR binds to the catalytic site of PPPs, this suggests that CYB5R4 has a selective effect on PP4 and PP6 catalytic subunits, potentially by affecting the catalytic site. To test if PP4 and PP6 activity was affected, we generated stable cell lines expressing affinity-tagged catalytic subunits of PP2A, PP4 and PP6 in U2OS parental and *CYB5R4* KO cells and performed peptide dephosphorylation assays with immunopurified catalytic subunits. This showed that PP4 and PP6 catalytic activity is impeded in the absence of CYB5R4, while PP2A activity is unaffected (**Fig. 1F**). The reduction of PP6 activity in the absence of CYB5R4 was confirmed in cells as we observed a strong increase in the PP6-specific target pT35 MOB1 in *CYB5R4* KO cells (**Fig. S1H**) (34). The loss of activity was not an effect of aberrant holoenzyme formation, as a comparison of immunopurified catalytic subunits from parental or *CYB5R4* knockout cells revealed no major defects in trimeric holoenzyme composition (**Fig. S1I-J, Table S7-8**).

In line with the evolutionary conservation of PPPs and their regulation, deletion of the *CYB5R4* orthologue *irc21* in *S*. *cerevisiae* resulted in depletion of Pph3 (PP4 C) and Sit4 (PP6 C) in PIB-MS (**Fig. S2A, Table S9**) and reduced catalytic activity of Pph3 and Sit4 but not Pph21 (PP2A C) *in vitro* (**Fig. 1G**). Additionally, the activity of Sit4 towards its substrate Sap185 was reduced *in vivo* without affecting holoenzyme composition (**Fig. S2B**). Our data show that Irc21 is a positive regulator of Sit4 and Pph3, and not Pph21, consistent with genetic data (35,36).

Intriguingly, Irc21 contains only a Cytb5 domain and not the CS or Cytb5-R domains (**Fig. 1D**), prompting us to ask whether the Cytb5 domain of CYB5R4 would be sufficient for its function in PP4/PP6 activation. Indeed, by re-introducing the domains separately in U2OS *CYB5R4* KO cells, we found that the Cytb5 domain was necessary and sufficient for rescuing the sensitivity to OA (**Fig. S2C-D**), arguing that at least in this cell culture assay, the CS and Cytb5-R domains are not required. Indeed, immunopurification of affinity-tagged CYB5R4^1-153^ followed by MS analysis revealed co-purification of PP4 and PP6 components, but not PP2A (**Fig. S2E, Table S10**). Gratifyingly, the human Cytb5 domain could complement the *irc21* deletion in yeast, rescuing the activity of Sit4 towards its substrate Sap185 *in vivo* (**Fig. S2F**).

Altogether, our genome-wide CRISPR screen uncovers PPP regulatory mechanisms and identifies CYB5R4 as an evolutionarily conserved activator of PP4 and PP6.

### Base editing tiling screen provides a high-resolution map of phosphatase function and regulation

To get deeper molecular insights into PPP regulation, we subjected established and candidate regulators and holoenzyme components scoring in the genome-wide OA synthetic lethality screen for base editor tiling screening. We selected a total of 19 targets choosing to include all PP2A-like catalytic subunits and known regulators in the library (**Fig. 2A-B**). Briefly, base editing relies on a Cas9 protein (often a Cas9 nickase) fused to a base editor being directed by a gRNA to a specific genomic locus where mutations are precisely introduced (**Fig. 2A**). The base editor makes mutations in an “editing window” at a specific distance from the protospacer adjacent motif (PAM), where the mutational outcome is determined by the underlying codons (37). We decided to use the efficient ABE8e adenine base editor, which makes A->G conversions in a 5 nucleotides editing window, fused to an SpG Cas9 nickase, which has a relaxed PAM requirement (NGN) that allows high mutational coverage of the protein-encoding sequence (38–42). We generated a custom 7928 gRNA ‘tiling’ library targeting the 19 genes, hereby mutating 5471 residues to cover 50.8% of the coding sequence (**Fig. 2B**). We used a similar screening setup as the CRISPR-Cas9 knockout synthetic lethality screen (**Fig. 1B**) with changes in gRNA abundance as a measure of mutational effect on cell fitness. Cellular fitness effects are interpreted in light of predicted amino acid mutations, allowing us to generate a functionally inferred residue map for each protein. By comparing gRNA abundance at endpoint of the screen (T_18_) to starting point (T_0_) as well as OA-treated (T_18_) to untreated (T_18_), we can map both residues whose mutation caused proliferation defects and residues whose mutation caused synthetic lethality with OA (**Fig. 2B, Fig. S3A**). This identified 585 and 378 gRNAs which caused proliferation defects and synthetic lethality in OA, respectively (using a z-score cut-off of <-5). These data are integrated with structural models of proteins and curated datasets, available to explore on an interactive website (https://slim.icr.ac.uk/base_editing/base_editing_tiling) to facilitate future research on PPP regulation.

**Figure 2:**
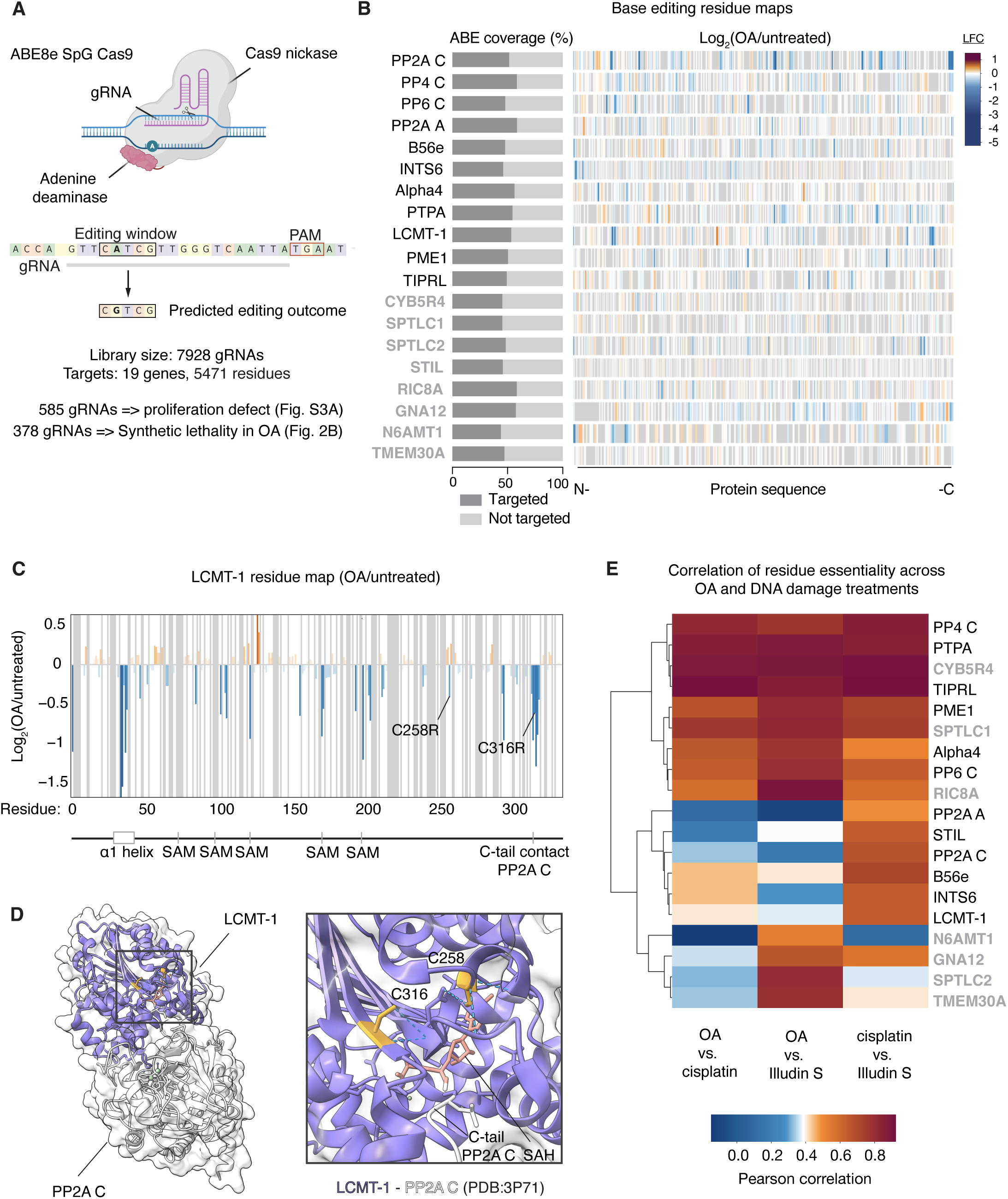
Base editing screen maps functional residues of PPP regulation. A) Schematic of adenine base editing with ABE8e-Cas9 SpG and information on base editing library. ABE, Adenine base editing. PAM, protospacer adjacent motif. B) Overview of synthetic lethality base editing screen results. Left: mutational coverage of the protein-coding sequence for each target. Right: residue maps of synthetic lethality in okadaic acid (OA) for each target. Each protein sequence is represented from left to right, and the color gradient represents the average log2 fold changes of gRNAs targeting the indicated residue in OA vs untreated conditions. Blue values specify that mutation of the target residue causes depletion (synthetic lethality) in OA and red enrichment. Grey specifies residues not targeted. Established PP2A-like holoenzyme components and regulators are shown in black and candidate regulators in grey. LFC, log2 fold change. C) Residue map of LCMT-1 showing synthetic lethality in OA. X-axis depicts the amino acid residue targeted for mutation and the y-axis the average log2 fold changes of gRNAs targeting the indicated residue in OA vs untreated conditions. Known functional entities of LCMT-1 are indicated. C258R and C316R are shown. SAM, S-adenosyl methionine. D) LCMT-1-PP2A structure (PDB:3P71) with annotation of C258 and C316. Hydrogen bridges are indicated in blue. SAH, S-adenosyl-homocysteine. E) Correlation analysis of residue essentiality per target across OA, cisplatin, and Illudin S treatments. Established components are indicated in black, candidate components are indicated in grey. Color scale represents Pearson correlation with blue indicating low correlation and red indicating high correlation.

The base editing approach was validated by the fact that mutations in residues known to be involved in activity, substrate specificity and regulation of PP2A-like phosphatases resulted in proliferation defects and synthetic lethality with OA (**Fig. 2B, Fig. S3A-B**). For example, residues coordinating the active site metal ions in the catalytic subunits and residues in B56epsilon involved in substrate binding scored in the screen (**Fig. S4A-D**). In addition to supporting known molecular mechanisms, the base editing maps also revealed possible novel mechanisms of regulation. As an example, mutation of two cysteine residues (C258 and C316) in proximity of C-tail contact residues in LCMT1 are synthetic lethal with OA, suggesting they could be part of novel regulatory mechanism tuning C-tail methylation involving disulfide bridge formation (**Fig. 2C-D**). Importantly, mutation of several residues amongst the novel regulators were synthetic lethal with OA (**Fig. S3B**), providing additional support for their function in PPP biology. For example, heme coordinating residues (H89/H112) in CYB5R4 (24), residues located at a binding pocket on SPTLC2 (R129 and I130) for the stimulatory protein ssSPTa (26,27), and multiple residues that are part of a binding interface between RIC8A (*e.g.*, S74 and L126) and the GNA12 C-terminal tail (N315, L321 and Q322) (43,44) scored in this screen (**Fig. 3A, Fig. S4E-H**).

**Figure 3:**
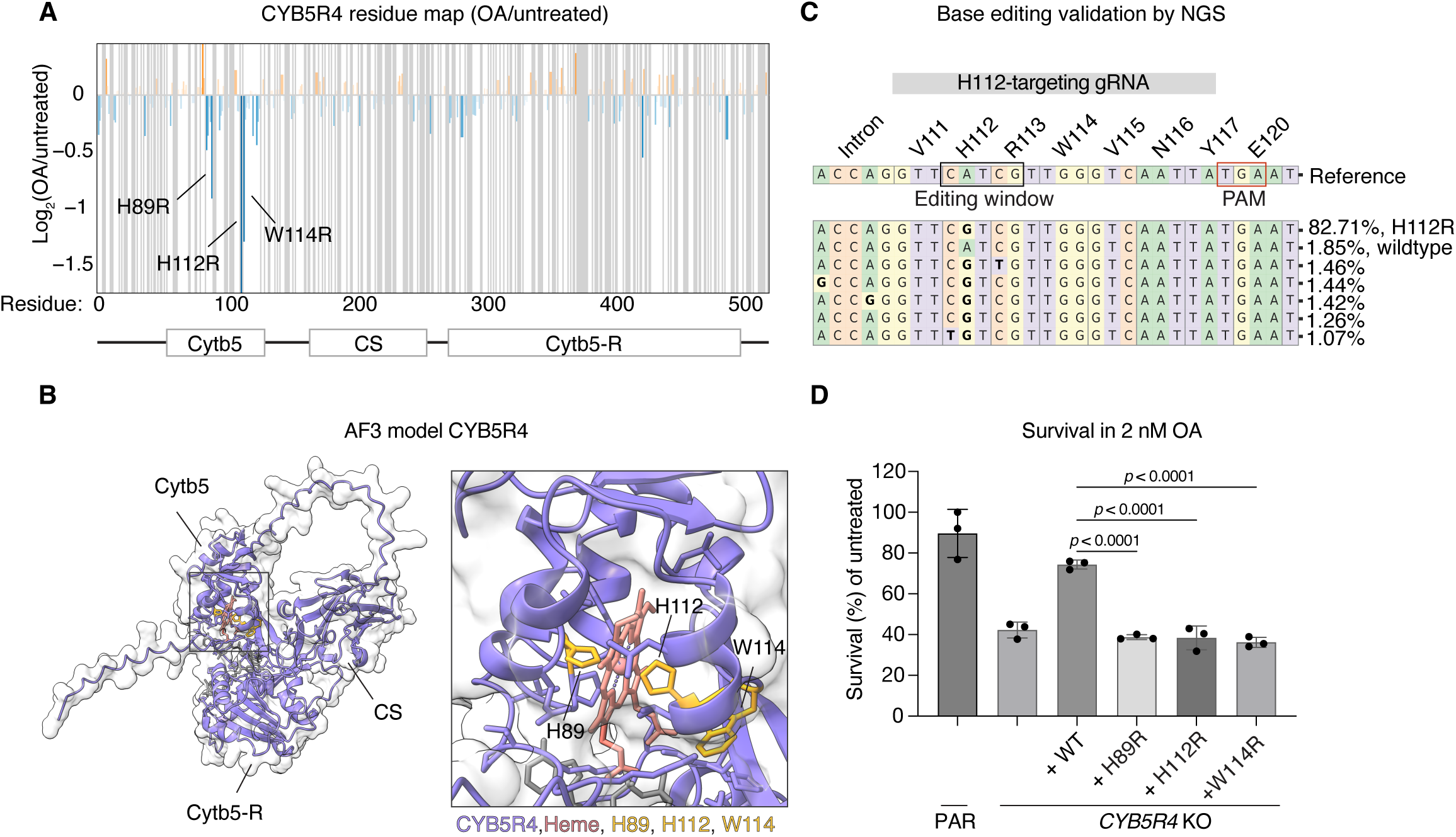
Cytb5 heme binding is essential for CYB5R4 function. A) Residue map of CYB5R4 showing synthetic lethality in OA. X-axis depicts the amino acid residue targeted for mutation and the y-axis the average log2 fold changes of gRNAs targeting the indicated residue in OA vs untreated conditions. The domains of CYB5R4 as well as H89R, H112R, and W114R are indicated. B) AlphaFold3 model of CYB5R4 with heme (salmon) and H89, H112, and W114 (yellow) indicated. C) Deep sequencing of the endogenous CYB5R4locus after transduction with a single gRNA targeting H112, followed by CRISPResso2 analysis shows the frequency of mutated alleles. Reference: the genomic sequence. The gRNA, PAM, and editing window are indicated as well as the amino acid translation. D) A colony formation assay was conducted in presence or absence of 2 nM OA with U2OS parental or *CYB5R4* knockout cells. Cells were stably complemented with full length CYB5R4-venus either wildtype (WT) or with the specified mutations. The survival is calculated as the relative number of colonies in OA to untreated and represents three independent experiments. Error bares depict standard deviations and the shown *p*-values are based on one-way ANOVA analysis with Tukey’s multiple comparisons test.

To determine whether our base editing tiling approach could be used to obtain new mechanistic insight into phosphatase function in intracellular signaling, we chose to probe the DNA damage response where PP2A-like phosphatases play a critical role (19,45). We thus generated functional residue maps in the presence of the DNA damaging agents cisplatin and Illudin S (**Fig. S5A-B**). Here, we identified 89 and 98 gRNAs which caused synthetic lethality with cisplatin and Illudin S, respectively, confirming that our screen can pick up phosphatase residues important for the response to DNA damaging agents. Correlating the mutational effects in OA, cisplatin, and Illudin S showed that PP4, PTPA, TIPRL, and CYB5R4 clustered together in all three treatments (**Fig. 2E**), arguing that mechanisms maintaining normal PP4 activity is key in determining response to the DNA damaging agents, consistent with known functions of PP4 (46,7). In contrast, PP2A C showed low correlation, revealing that loss of PP2A function does not further sensitize cells to these DNA damaging agents.

### The heme group is required for CYB5R4 function and binding to PP4/PP6

We were intruiged by the observation that heme coordinating residues (H89/H112) in CYB5R4 as well as a proximal trypthophan residue (W114) scored in our base editor screen (**Fig. 3A-B**), suggesting a key role of the heme group in activating PP4 and PP6. To pursue these observations further, we first established that the gRNAs introduced the predicted mutations. We performed CRISPResso2 mutational analysis of the target loci by deep sequencing in RPE-1 cells transduced with the single gRNAs. Indeed, the analysis showed efficient introduction of the predicted mutations targeting H89 (Y88 was also frequently targeted), H112, and W114 (**Fig. 3C, Fig. S5C**). Next, to validate the screen results, we re-introduced CYB5R4 carrying the single base editing mutations in U2OS *CYB5R4* knockout cells (**Fig. S5D**), which revealed that these mutations conferred hypersensitivity to OA and cisplatin (**Fig. 3D, Fig. S5E-G**).

Since the heme-coordinating histidines and the proximal tryptophan are important for CYB5R4 function, we asked whether they would be required for the observed binding to PP4/6 (**Fig. S2E**). Indeed, when either H89/H112 or W114 were mutated to alanine, binding of PP4 and PP6 components to CYB5R4^1-153^ was lost (**Fig. 4A, Table S10**). Additionally, we observed that all the catalytic subunits of the calcineurin phosphatase specifically co-purified with CYB5R4^1-153^ wild-type (**Table S10**), consistent with previous results (47), suggesting that CYB5R4 may function beyond the PP2A-like phosphatases.

**Figure 4:**
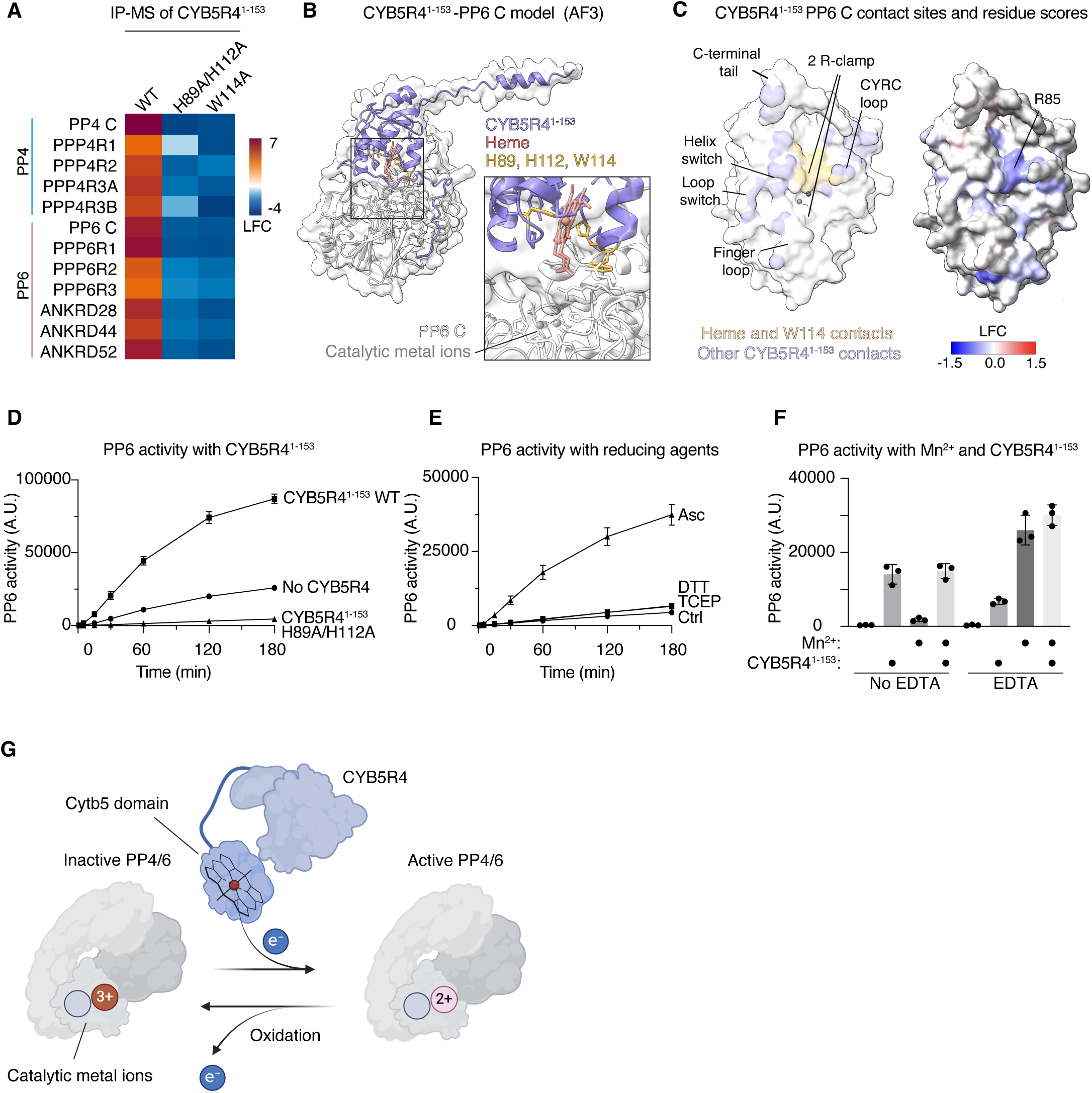
CYB5R4 activates PP4 and PP6 by reducing the metal ions in the catalytic site. A) Heatmap comparing the interactomes of venus-tagged CYB5R4^1-153^ wildtype (WT), H89A/H112A, and W114A to venus control, which were immunoprecipitated from HeLa cells and analysed by mass spectrometry. PP4/6 holoenzyme components are shown. Colors represent log2 fold changes with red being increased and blue depleted compared to venus control. B) AlphaFold3 model of CYB5R4^1-153^ and PP6 C with heme and residues H89, H112 and W114 indicated. The N-terminal tail of CYB5R4 is not shown in the close-up views for clarity. C) AlphaFold3 model of PP6 with annotation of CYB5R4^1-153^ contacts (left) and a color gradient (right) that represents the average log2 fold changes of gRNAs targeting the indicated residue in OA vs untreated conditions. Blue values specify that mutation of the target residue causes depletion (synthetic lethality) in OA and red enrichment. Grey specifies residues not targeted. D) DiFMUP dephosphorylation assay measuring the activity of purified PP6 holoenzyme in presence or absence of pre-reduced purified CYB5R4^1-153^ WT or H89A/H112A. Data is from three independent experiments, and error bars represent standard deviations. E) DiFMUP dephosphorylation assay measuring the activity of purified PP6 holoenzyme in presence or absence of the indicated reducing agents at 1 mM. Data is from three independent experiments, and error bars represent standard deviations. F) DiFMUP dephosphorylation assay measuring the activity of purified PP6 holoenzyme, which was first pre-incubated with or without EDTA to extract metal ions. Next, the enzyme was incubated in presence or absence of Mn^2+^ and finally in the presence or absence of pre-reduced purified CYB5R4^1-153^ WT. Activity after 30 minutes is shown (also see Supplementary Figure S7F). Data is from three independent experiments, and error bars represent standard deviations. G) Model of PP4/6 regulation by CYB5R4.

Given that the regulatory and scaffold subunits of PP4 and PP6 are very different in sequence, but their catalytic subunits are strikingly similar, we reasoned that CYB5R4 would bind to the catalytic subunits. We therefore generated AlphaFold 3 (AF3) models of CYB5R4^1-153^ and the catalytic subunits of PP4 and PP6 to obtain further insights into their mode of interaction (**Fig. 4B, Fig. S6A-B**) (48). In line with our hypothesis, the AF3 models confidently predicted interaction of the Cytb5 domain with the phosphatase catalytic subunits. More specifically, the Cytb5 domain was positioned facing the catalytic site, in such a way that the heme group and W114 closely contacted residues on PP4 (Y125, C263) and PP6 (R85) whose mutations were synthetic lethal with OA (**Fig. 4C, Fig. S6C**). This AF3 model explains the requirement for heme coordination by H89/H112 and W114 for binding to the catalytic subunit and suggests that CYB5R4 competes with other regulators, such as PTPA, for binding to the catalytic subunit (10). Indeed, in the absence of CYB5R4/Irc21, we observe increased binding of PTPA/Rrd1 to the catalytic subunits, potentially reflecting this competition (**Fig. S1I, Fig. S6D**).

### CYB5R4 activates PP4 and PP6 by reducing the metal ions in the catalytic site

The positioning of the CYB5R4 heme group in close proximity to the catalytic site of PP4 and PP6 in AF3 models suggested to us that CYB5R4 activates these phosphatases by donating electrons from the heme group to their catalytic site. This could be facilitated by a relay of electrons from the heme group through W114 (23,24). To test this directly, we established an *in vitro* reconstituted system with purified components. To this end, we incubated purified reconstituted PP4 and PP6 holoenzymes, expressed with the Multi-Mam system in HEK293, with purified recombinant CYB5R4^1-153^ wild-type or H89A/H112A and tested their activity. While PP4 and PP6 both displayed modest baseline activities, CYB5R4^1-153^ wild-type stimulated their activities over 3-fold, whereas the H89A/H112A mutant was unable to do so (**Fig. 4D, Fig. S7A**). These findings were fully recapitulated in yeast, where inactive Sit4 (PP6) isolated from the irc21 deletion strain could be rescued by human CYB5R4^1-153^ wildtype but not H89A/H112A *in vitro* (**Fig. S7B-D**). Importantly, in each of these assays, only CYB5R4^1-153^ pre-reduced by L-ascorbate could induce activation, corroborating our hypothesis that CYB5R4 acts by donating electrons to PP4 and PP6. The most likely targets for reduction by CYB5R4 would be either the catalytic metal ions or cysteine residues, since both are prone to oxidation (49,50). However, in our *in vitro* assays, PP4 and PP6 activity was not stimulated by the cysteine reducing agents (DTT or TCEP), while L-ascorbate that can reduce metal ions stimulated phosphatase activity (**Fig. 4E, Fig. S7E**). This argues that in our *in vitro* assays, CYB5R4 stimulation is through metal ion reduction. To support this hypothesis, we converted PP6 into an oxidation-resistant phosphatase by removing the active site metal ions by incubation with EDTA and re-activated the phosphatase by adding back bivalent metal ions in the form of Mn^2+^. The activity of this PP6-Mn^2+^ complex was not further stimulated by CYB5R4 (**Fig. 4F, Fig. S7F**). Together, these observations support the role of CYB5R4 as a reductase for the active site metal ions (**Fig. 4G**).

## Discussion

Here, we integrate CRISPR knockout and base editor screens to uncover novel molecular mechanisms of phosphatase regulation - a general approach that can be applied to other signaling molecules where specific inhibitors are available. We provide a readily accessible web resource for consulting our high-resolution functional map of PP2A-like phosphatase regulators and holoenzyme components, facilitating future research on mechanisms controlling their activity. Our map can be used to broaden our understanding of cross talk between PP2A-like phosphatases and intracellular signaling mechanisms, as exemplified by CYB5R4, which we pursued in depth. We identified the protein as an evolutionary conserved reductase for the active site metal ions, uncovering crosstalk between the redox state of the cell and PP4 and PP6 activity. This expands the regulatory mechanisms of PPPs and suggests that these phosphatases likely share regulatory principles with protein tyrosine phosphatases, where dedicated enzymes controlling active site oxidation and reduction is a central and well-established mechanism (51–53). The presence of such a mechanism fits with the fact that PPPs have been shown to be inactivated through catalytic metal ion oxidation (49,54). Whether a dedicated oxidase for the active site of PP4 and PP6 exists, as reported for PP1 (55), or whether the active site metal ions are highly sensitive to oxidation is presently unclear. While the exact nature of these metal ions is debated, Fe^2+^ has been proposed, and importantly, this ion is readily oxidized (3,49,56). An intriguing observation is that PP2A activity is not dependent on CYB5R4 function, despite a high level of sequence identity with PP4 and PP6. Whether this reflects a difference in the nature of active site metal ions, or subtle differences in the active site architecture that make the metals more sensitive to oxidation in PP4 and PP6 is unclear. Our results argue that PP4 and PP6 activity respond to the redox state of the cell through metal ion oxidation while PP2A activity does not, revealing a fundamental difference in their regulation.

## Materials and Methods

### Cell culture

Cells (RPE-1;hTERT;TP53-/-;Cas9, RPE-1;hTERT;TP53-/-, U2OS Flp-In T-REx (referred to as U2OS), and HeLa) were cultured in Dulbecco’s Modified Eagle Medium with GlutaMAX (Gibco) supplemented with 10 % fetal bovine serum (Gibco) and 10 units/mL of penicillin and 10 μg/mL of streptomycin (Gibco) at 37°C with 5% CO_2_. Low oxygen experiments with U2OS cells were performed in an oxygen control chamber (InvivO_2_ 400 Hypoxia workstation, Baker Ruskinn) set at 37°C, 5% CO_2_ and 6% O_2_. Cell lines used in this study are listed in Table S11.

### CRISPR-Cas9 KO screens

#### Virus production

Lentiviral particles were produced by co-transfection of the sgRNA plasmid library TKOv3::pLCKO2, with lentiviral packaging plasmids pMD2.G and psPAX2 in HEK293T/17 cells (ATCC, CRL-11268) using Lipofectamine 3000 (Invitrogen) in Opti-MEM (Gibco). 6 hours after transfection, medium was exchanged for DMEM GlutaMax + 10% FBS + 100 U/mL penicillin–streptomycin + 1% bovine serum albumin. 48 hours after transfection, viral particles were harvested and filtered through a 0.45 μm syringe filter before freezing at −80°C. The TKOv3::pLCKO2 library (Addgene plasmid #125517) was a gift from Jason Mofat. pMD2.G (Addgene plasmid # 12259; http://n2t.net/addgene:12259; RRID:Addgene_12259) and psPAX2 (Addgene plasmid # 12260; http://n2t.net/addgene:12260; RRID:Addgene_12260) were gifts from Didier Trono.

#### Transduction and cell culture

RPE1;hTERT;*TP53*-/-;Cas9 cells (a kind gift from D. Durocher) were cultured in DMEM GlutaMax supplemented with 10% FBS and 100 U/mL penicillin–streptomycin and passaged every three days. The screen was performed as a duplicate (1 single transduction split up in 2 replicate drug treatments) at a coverage of above 300-fold sgRNA representation, which was maintained throughout the screen. Cells were transduced with the lentiviral library at a low multiplicity of infection (0.2-0.3) by treating cells with 8 μg/mL polybrene and lentiviral supernatants for 24 hours. Transduced cells were selected by treatment with 20 μg/mL puromycin for 24 hours followed by trypsinization and reseeding in the same plates with 20 μg/mL puromycin for another 24 hours. After selection, cells were passaged for 6 days before splitting into untreated or okadaic acid (OA) treated fractions. Cells were passaged for an additional 12 days with passaging every 3 days in medium with or without a low dose of OA (2 nM), which corresponds to predetermined LD_20_ concentrations. Cell pellets were harvested after completion of selection, which we consider the start of the screen, (T_0_) and at the final timepoint (T_18_).

#### Next generation sequencing and analysis

Genomic DNA was extracted from the cells, and the genomic DNA regions containing the integrated sgRNAs were amplified by PCR using NEBNext Ultra II Q5 Master mix (New England BioLabs) with the pLCKO2_forward and pLCKO2_reverse primers (Table S12). A second PCR reaction introduced i5 and i7 multiplexing barcodes (Table S12) and gel-purified PCR products were sequenced on Illumina NextSeq500. Data was analysed as in (57). Briefly, fastq files were generated using bcl2fastq v2.20.1, reads were trimmed to 20 bp using cutadapt 1.18 (58) and trimmed reads were assigned to guides in the TKOv3 library by MAGeCK 0.5.9.5 (59) to create a count matrix, from which gene scores (NormZ) were calculated with DrugZ (60). Data quality was assessed by BAGEL.py “pr” function (61) with core essential and non-essential gene lists (https://github.com/hart-lab/bagel), comparing T_0_ to T_18_ of untreated cells.

### Generation of U2OS Flp-In T-REx *CYB5R4* knockout cells

A gRNA targeting CYB5R4 (AATTGACCCAACGATGAAC; (19)) was synthesized as DNA oligonucleotides with overhangs for BbsI cloning: Forward *CACCG*AATTGACCCAACGATGAACC; Reverse: *AAAC*GGTTCATCGTTGGGTCAATT*C* (cloning sites in italic). The oligos were annealed and cloned into pSpCas9(BB)-2A-Puro (pX459) using BbsI cloning (62). (pSpCas9(BB)-2A-Puro (pX459) was a gift from Feng Zhang (Addgene plasmid # 48139; http://n2t.net/addgene:48139; RRID:Addgene_48139). U2OS Flp-In T-REx cells (a kind gift from H. Piwnica-Worms) were transfected with the gRNA encoding plasmid using JETOptimus (Polyplus) and selected for 48 hours in 1 μg/mL puromycin followed by growth until colonies formed. Single colonies were picked, expanded in medum supplemented with 5 μg/mL blasticidin S HCl (Sigma) and 100 μg/mL Zeocin (Invitrogen), and *CYB5R4* knockout was verified by Western blot analysis.

### Cloning of plasmids

The human CYB5R4 cDNA was obtained from Promega and cloned into pcDNA5/FRT/TO/ plasmids with N- or C-terminal venus- or myc-tagging using restriction enzyme cloning. Mutations were introduced by mutagenic PCR with the primers indicated in in Table S12. Fragments of CYB5R4 for domain analysis were cloned by PCR with restriction site overhangs and subsequent restriction enzyme cloning. PP2A, PP4, and PP6 catalytic subunits were cloned into pcDNA5/FRT/TO with 3xFLAG-tagging using restriction enzyme cloning. All primers and plasmids used in this study are listed in Table S12 and S13, respectively.

### Generation of stable U2OS Flp-In T-REx cells

U2OS Flp-In T-REx cells or U2OS Flp-In T-REx *CYB5R4* knockout cells were grown in medium supplemented with 5 μg/mL blasticidin S HCl (Sigma) and 100 μg/mL Zeocin (Invitrogen). To generate stable cell lines in the Flp-In system, cells were co-transfected with pOG44 and a pcDNA5 plasmid encoding the indicated construct (Table S13) in a 10:1 ratio using the JETOptimus transfection reagent. After transfection, stable Flp-In T-REx cells were selected in medium supplemented with 5 μg/mL blasticidin S HCl and 200 μg/mL Hygromycin B. Expression from the Tet-ON inducible promoter in Flp-In T-REx cells was induced with the doxycycline doses as indicated in Table S11. In U2OS Flp-In T-REx *CYB5R4* knockout cells, the doxycycline doses required to achieve expression levels of exogenous CYB5R4 constructs equal to or higher than endogenous CYB5R4 were estimated by Western blot analysis.

### WB analysis of cell extracts

Cells were harvested by trypsinization and lysed on ice in RIPA buffer (10 mM Tris, pH 7.4, 150 mM NaCl, 1 mM EDTA, 1% NP-40, 0.5% sodium deoxycholate, 0.1% SDS) supplemented with 1 mM DTT and Complete protease inhibitor cocktail (Roche). The lysate was cleared by centrifugation at 20,000 g at 4°C for 30 minutes, and BCA assay (Pierce) was used to even out protein concentrations between samples. Samples were analyzed by SDS-PAGE and Western blotting using the antibodies indicated in Table S14. For WB analysis with phospho-antibodies, RIPA buffer was additionally supplemented with PhosSTOP phosphatase inhibitor cocktail (Roche), and lysates were sonicated with Bioruptor Plus (Diagenode) prior to clearing.

### PIB-MS

Cells (quadruplicate for each condition) were harvested by trypsinization, pelleted, snap frozen and processed as previously described (33). Proteins were enriched from eluates using the SP3 method (63) and digested overnight in 25 mM ammonium bicarbonate with trypsin for mass spectrometric analysis. Digests were analyzed using either a Q-Exactive Plus quadrupole Orbitrap mass spectrometer (ThermoScientific) equipped with an Easy-nLC 1000 (ThermoScientific) or an Orbitrap Fusion Lumos mass spectrometer (ThermoScientific) equipped with an Easy-nLC 1200 (ThermoScientific), and nanospray source (ThermoScientific). COMET (release version 2014.01) in high-resolution mode was used to search raw data (64) against a target-decoy (reversed) (65) version of the human proteome sequence database (UniProt; downloaded 8/2020), mouse proteome sequence database (UniProt; downloaded 8/2020), or *Saccharomyces cerevisiae* proteome sequence database (UniProt; downloaded 8/2020) with a precursor mass tolerance of ± 1 Da and a fragment ion mass tolerance of 0.02 D requiring fully tryptic peptides (K, R; not preceding P) and up to three mis-cleavages. Static modifications included carbamidomethylcysteine and variable modifications included oxidized methionine. Searches were filtered to a < 1% FDR at the peptide level. Quantification of LC-MS/MS spectra was performed using MassChroQ (66) and the iBAQ method (67). Missing PPP subunit abundances were imputed and normalized across all samples by quantile normalization in Perseus (68). Statistical analysis was carried out by a two-tailed Student’s t-test, and heatmaps were generated in Perseus.

### IP-MS

Doxycycline-inducible 3xFLAG, 3xFLAG-PP2A-catalytic (C), 3xFLAG-PP4-C and 3xFLAG-PP6-C U2OS Flp-In T-Rex wild-type and *CYB5R4* KO stable cell lines (generated as described above) were treated with 10 ng/ml doxycycline. HeLa cells were transiently transfected using JetOptimus with 2 μg of venus, CYB5R4^1-153^-venus, CYB5R4^1-153^-venus H89A/H112A and CYB5R4^1-153^-venus W114A. 24 hours after doxycycline induction or transfection, cells were washed in PBS and lysed in low salt lysis buffer (50 mM NaCl_2_, 50 mM Tris.HCl pH 7.4) supplemented with 0.1% NP-40 and cOmplete protease inhibitor cocktail (Roche) followed by sonication using the Bioruptor Plus at 4°C. Lysates were cleared by centrifugation for 30 minutes at 14,000 g whereafter supernatants were incubated with Fab-trap beads (Proteintech) for immunoprecipitation of 3xFLAG control and 3xFLAG-conjugated catalytic subunits or with GFP-trap beads (Proteintech) for immunoprecipitation of venus-tagged CYB5R4 wild-type and mutant. After rotating at 4°C for 1 hour, beads were washed 3 times in low salt lysis buffer (2 times with 0.1% NP-40, 1 time without) and once in TBS. Dry beads were subsequently resuspended in 2X Laemli sample buffer and boiled at 95°C for 10 minutes. Proteins were enriched from eluates using the SP3 method (63) and digested overnight in 25 mM ammonium bicarbonate with trypsin for mass spectrometric analysis. Digests were analyzed using an Orbitrap Fusion Lumos mass spectrometer (ThermoScientific) equipped with an Easy-nLC 1200 (ThermoScientific), and nanospray source (ThermoScientific). COMET (release version 2014.01) in high-resolution mode was used to search raw data (64) against a target-decoy (reversed) (65) version of the human proteome sequence database (UniProt; downloaded 8/2020) with a precursor mass tolerance of ± 1 Da and a fragment ion mass tolerance of 0.02 D requiring fully tryptic peptides (K, R; not preceding P) and up to three mis-cleavages. Static modifications included carbamidomethylcysteine and variable modifications included oxidized methionine. Searches were filtered to a < 1% FDR at the peptide level. Quantification of LC-MS/MS spectra was performed using MassChroQ (66) and the iBAQ method (67). Missing protein abundances were imputed and bait abundances were normalized across all samples (68) Statistical analysis was carried out by a two-tailed Student’s t-test, and heatmaps were generated in Perseus.

### Proteome analysis of RPE-1 wild-type and CYB5R4 KO cells

Cells (quadruplicate for each condition) were harvested by trypsinization, pelleted, snap frozen and processed as previously described (33). Proteins were using 5 mM DTT and 15 mM iodoacetamide and iodoacetamide, respectively. Samples were incubated overnight at 37°C with 1:100 (w/w) trypsin. The next day, the trypsin digest was stopped by the addition of 0.25% TFA (final v/v). Precipitated lipids were removed by centrifugation (3500 x g for 15 minutes), and the peptides in the supernatant were desalted over an Oasis HLB plate (Waters). Peptides were labeled with Tandem-Mass-Tag (TMT) reagent (Thermo Fisher Scientific). Once labeling efficiency was confirmed to be at least 95%, each reaction was quenched by the addition of hydroxylamine to a final concentration of 0.25% for 10 minutes, mixed, acidified with TFA to a pH of about 2, and desalted over an Oasis HLB plate (Waters). The desalted multiplex was dried by vacuum centrifugation and separated by offline Pentafluorophenyl (PFP)-based reversed-phase HPLC fractionation as previously described (69). TMT-labeled peptides were analyzed on an Orbitrap Lumos mass spectrometer (Thermo Scientific) equipped with an Easy-nLC 1200 (Thermo Scientific) and nanospray source (Thermo Scientific). Raw data was searched and processed as previously described (70). Peptide intensities were adjusted based on total TMT reporter ion intensity in each channel, and log_2_ transformed. P-values were calculated using a two-tailed Student’s t-test.

#### Activity assays with phosphatases immunopurified from cells

Immunoprecipitation of was performed similar as for IP-MS but differing at lysis and washing steps. Specifically, for lysis samples were left on ice for 20 minutes with vortexing every 5 minutes and washes were performed 2 times with low salt lysis buffer (50 mM NaCl_2_, 50 mM Tris.HCl pH 7.4) supplemented with 0.1% NP-40 and one time in activity assay buffer (50 mM NaCl_2_, 150 mM Tris.HCl pH 7.4). Subsequently, dry beads were resuspended in activity assay buffer and divided across conditions. Phosphopeptide WRRA(pT)VA (Peptide2.0) dissolved at 1mM in 100 μM Tris.HCl pH 8 was added to a final concentration of 230 μM and dephosphorylation reactions were incubated at 30°C. Reactions were stopped at indicated timepoints by adding to malachite green (PicolorLock, Abcam), and activity was determined by means of absorbance measurements at 620 nm in a multi-well plate reader (Fluostar Omega, BMG Labtech). Input levels of immunopurified phosphatases were determined by western blot. Activities were corrected for background using absorbance measured in 3xFLAG control conditions. Experiments were performed in triplicate. Data was analyzed in PRISM10 (Graphpad).

### Colony formation assays

The cells were trypsinized, resuspended in medium, and counted. 200 cells were seeded per well in 6-well plates in minimum two wells per condition with the doxycycline concentration indicated in Table S11. 24 hours after seeding, cells were treated with the indicated compounds (2 nM okadaic acid or 1 μM cisplatin) or left untreated. After 7 additional days of growth, formed colonies were fixed and stained in a methyl violet solution (0.5% methylviolet, 25% methanol), and the number of colonies was quantified on a GelCount (Oxford optronix). The survival after treatment with a given compound is calculated as the number of colonies after treatment normalized to the number of colonies of the untreated condition. All experiments were performed as biologically independent triplicates and analyzed in PRISM10 (Graphpad). One-way ANOVA analysis with Tukey’s multiple comparisons tests were performed to test for statistical significance.

### Protein purification

#### CYB5R4 cloning, expression, and purification

CYB5R41-153 was cloned into pCPR0063 using LIC cloning allowing expression of a His-GST-TEV fusion protein. The protein was expressed overnight in BL21(DE3) at 18°C and harvested by centrifugation. Cell pellets where resuspended in lysis buffer (50 mM NaP pH=7.5, 300 mM NaCl, 10 mM imidazole, 10% glycerol, 0,5 mM EDTA, Benzonase, and protease inhibitor cocktail). The cells where lysed by sonication and cleared by centrifugation and the supernatant applied to a 5 ml HiTrap Nickel column and washed with run buffer (50 mM NaP pH=7.5, 300 mM NaCl, 10 mM imidazole, 10% glycerol, 0,5 mM TCEP) and protein eluted with a 10-500 mM imidazole gradient (100 CVs). The protein was dialyzed and cleaved with TEV and applied to a 5ml HiTrap Nickel column to remove His-GST. The flowthrough was concentrated and applied to a Superdex 75 16/60 column equilibrated with 50 mM NaP pH=7.5, 150 mM NaCl, 10% glycerol, 0,5 mM TCEP, and peak fractions pooled and stored at −80°C.

#### PP4 and PP6 holoenzyme cloning, expression, and purification

The PP4 and PP6 holoenzyme expression constructs were generated following the ACEMBL Multimam guidelines. Briefly, PPP4R3A and PPP6R1 where cloned into pACEMam1 and the PPP4R2/FLAG-PPP4C and ANKRD28/FLAG-PPP6C into the donor vectors pMDC and pMDK. A 2-step recombineering protocol was followed to obtain the final pACEMam1 constructs with all 3 cDNAs. HEK293 cells were grown in FreeStyle F17 complete medium and transfected at a cell density of 1×10E6. Cells were transfected using OPTIMEM and PEI using 1μg DNA per 1 ml of culture and following standard transfection procedures. Cells were harvested after 48 hours and processed for purification of complexes. Pellets were lysed in lysis buffer (75 mM Tris-HCl PH 7.5, 10% glycerol, 2mM MgCl2, 150 mM NaCl, 0.1⁒ w/v Tween 20, 0.008 ⁒ w/v NP-40) supplemented with complete protease inhibitor cocktails (Roche) and 8 U/mL Benzonase nuclease (Millipore) using pressure homogenizer EmulsiFlex-C3 (Avestin). The lysates were cleared at 14400 rpm at 4°C for 1 h and again for another 30 min. Pre-equilibrated anti-FLAG M2 affinity gel (Sigma Aldrich) was added to the supernatant incubated 1h at 4°C with rotation. The beads were spun down and washed first in FLAG buffer (75 mM Tris-HCl PH 7.5, 10% glycerol, 2mM MgCl2, 150 mM NaCl), then in FLAG buffer supplemented with 5 mM ATP Mg, and then again in FLAG buffer. The protein was eluted in FLAG buffer supplemented with 0.4 mg/mL 3xFLAG peptide (Sigma Aldrich). Protein complexes were analyzed by analytical size exclusion chromatography.

### Activity assays with recombinant proteins

Activity assays were performed in activity assay buffer (150 mM NaCl, 75 mM Tris.HCl pH 7.4) by mixing purified PP4 and PP6 holoenzymes (in FLAG buffer, 150 mM NaCl, 75 mM Tris.HCl pH 7.4, 2 mM MgCl_2_, 10% glycerol) with reaction components (below), transferring the mix to 96-well plates, and adding 6,8-Difluoro-4-Methylumbelliferyl Phosphate (DiFMUP, Invitrogen) to a final concentration of 100 μM. Baseline activity was determined immediately after adding DiFMUP by measuring fluorescence at 350 nm excitation/450 nm emission in a microplate reader (Fluostar Omega, BMG Labtech), followed by incubation at 30°C with fluorescence sampling at 355 nm excitation/460 nm emission performed at indicated times. For each timepoint the baseline activity was subtracted. For CYB5R4 reactivation assays, CYB5R4^1-153^ WT, H89A/H112 (final concentration approx. 100 μg/μl), or FLAG buffer control was pre-reduced with 1 mM L-ascorbate for 15 minutes at room temperature before addition to the reaction. For phosphatase activation with reducing reagents, phosphatases were incubated with 1 mM DTT, TCEP and L-ascorbate for 15 minutes at room temperature before DiFMUP addition. For EDTA treatments, PP6 holoenzyme purified in FLAG buffer without MgCl_2_ was incubated with 15 μM EDTA at 30°C overnight in activity buffer supplemented with 1 mM DTT to prevent cysteine oxidation. This was followed by addition of metal ion (MgCl_2_) to a final concentration of 50 μM in activity buffer and incubation for 10 min at room temperature. Finally, pre-reduced CYB5R4^1-153^ WT or FLAG buffer control was added before DiFMUP addition.

### Animal experiments

#### Animal husbandry

Global Ncb5or-null (KO) mouse was generated through genetic ablation of exon 4 (30) and backcrossed into C57BL/6J for >12 generations prior to this study. The KO and wild-type (WT) embryos (littermates) were produced from heterozygous crossing in a pathogen-free facility at 24°C under a 12-hour light cycle with unlimited access to water and a standard rodent chow. All animal experiments were performed in accordance with the National Institutes of Health Guide for the Care and Use of Laboratory Animals and approved by the University of Kansas Medical Center Institutional Animal Care and Use Committee.

#### Harvest and culture of mouse embryonic fibroblasts (MEFs)

MEFs (E14.5) were collected from 14-day post-coitum pregnant dams (https://app.jove.com/v/3854/preparation-mouse-embryonic-fibroblast-cells-suitable-for-culturing). Immediately after each dam was anesthetized and cervical dislocated, the uterine horns were removed and placed in a clean 10-cm petri dish and washed three times with sterile phosphate-buffered saline (PBS, no Ca^2+^/Mg^2+^). Embryos were released into sterile PBS and processed individually under sterile conditions. Upon the removal of tail for genotyping and visceral (red or dark) tissues for disposal, each embryo was washed with 10 ml PBS three times, minced thoroughly with a curved iris scissor in a total of 7 ml Trypsin/EDTA digestion solution, and incubated at 37°C for ∼ 20 minutes with repeated pipetting until few chunks remained. After an additional 10-minute incubation, each digestion mixture was neutralized with ∼20 ml culture medium (see below for composition) and transferred to a 50 ml conical tube for thorough mixing. The content was then evenly added to a T25 culture flask containing 5 ml prewarmed culture medium, placed in a 37°C incubator overnight, and changed to fresh medium the next day to remove cell debris. When each flask became 80-90% confluent, cells were transferred to new T75 flasks for further expansion (1:5 to 1:3 for each passage). After passage 1 or 2, cell stocks were prepared from each MEF line in cryopreservation medium (each vial equivalent to one T25 flask) for long-term storage in liquid nitrogen or expedited shipment on dry ice for additional analyses. Culture medium: DMEM with 4.5 g/L glucose, L-glutamine, sodium pyruvate, plus 10% fetal bovine serum (FBS, heat-inactivated), 1% non-essential amino acids, and 1% Penicillin/Streptomycin. Cryopreservation medium: 90% FBS, 10% dimethyl sulfoxide (DMSO).

#### Genotyping

Polymerase chain reaction (PCR) was performed on genomic DNA of all embryos or adult breeders. Tissues were digested at 55°C overnight in lysis buffer (100 mM Tris-HCl, pH 8.8; 5 mM EDTA; 0.2% SDS; 200 mM NaCl; 0.1 mg/ml proteinase K added immediately before use). After a 5-minute high-speed centrifugation, the clear supernatant was diluted 10x with sterile water and directly subjected to genotyping PCR as previously described (30). Primer sequences and PCR conditions available upon request.

### CRISPR base editing tiling screen

#### Library design and cloning

A set of guide RNAs (gRNA) was designed to tile missense mutations across a selected set of 19 proteins (listed in Figure 2B) using the SpG Cas9 ABE8e adenine base editor (40). To predict the mutations resulting from ABE edits, we assumed full editing efficiency within the “editing window” of nucleotides 4–8 of the gRNA-targeted DNA sequence. Only gRNAs predicted to introduce amino acid substitutions were retained for further analysis. Positive (essential splice sites) and negative (non-targeting and intergenic) control gRNAs were also included to benchmark the screen. The gRNA library was synthesized (GenScript) as an oligonucleotide pool that followed a previously published design for amplification and Esp3I cloning (40) : 5’-[Forward primer (20 nt)]CGTCTCACACCG[sgRNA (20 nt)]GTTTCGAGACG[Reverse Primer (20 nt)]. The gRNA oligonucleotide pool was amplified using NEBNext Ultra II Q5 Master Mix (New England BioLabs) and the primers: Forward: GTGTAACCCGTAGGGCACCT; Reverse: GTCGAGAGCAGTCCTTCGAC, and then cloned into the Abe8e-Cas9-SpG lentiviral vector pRDA_479 (40) using Golden gate cloning with Esp3I and T7 ligase. pRDA_479 was a gift from John Doench & David Root (Addgene plasmid # 179099; http://n2t.net/addgene:179099; RRID:Addgene_179099). The plasmid library was purified first by PCR purification (NucleoSpin Gel and PCR Clean-up, Macherey-Nagel) and then isopropanol precipitation. Purified plasmid library was electroporated into Endura Electrocompetent cells (Lucigen), which were grown at 30 °C for 16 h on agar with 100 μg/mL carbenicillin. The colonies were scraped of the plates and plasmid DNA prepared (NucleoBond Xtra Maxi, Macherey-Nagel). The library was sequenced to confirm gRNA representation. This was done by PCR amplification from the library using NEBNext Ultra II Q5 Master mix and primers from Table S12, gel purification of amplicons, and finally sequencing on a NextSeq2000 (Illumina).

#### Virus production

Lentiviral particles were produced by co-transfection of the sgRNA plasmid library with lentiviral packaging plasmids pMD2.G and psPAX2 in HEK293T/17 cells (ATCC, CRL-11268) using Lipofectamine 3000 (Invitrogen) in Opti-MEM (Gibco). pMD2.G (Addgene plasmid # 12259; http://n2t.net/addgene:12259; RRID:Addgene_12259) and psPAX2 (Addgene plasmid # 12260; http://n2t.net/addgene:12260; RRID:Addgene_12260) were gifts from Didier Trono. 6 hours after transfection, medium was exchanged for DMEM GlutaMax + 10% FBS + 100 U/mL penicillin– streptomycin + 1% bovine serum albumin. 48 hours after transfection, viral particles were harvested and filtered through a 0.45 μm syringe filter before freezing at −80°C.

#### Transduction and cell culture

RPE1;hTERT;*TP53*-/-cells (a kind gift from D. Durocher) were cultured in DMEM GlutaMax supplemented with 10% FBS and 100 U/mL penicillin–streptomycin and passaged every three days. The screen was performed as a duplicate (two separate transductions) at a coverage of above 500-fold sgRNA representation, which was maintained throughout the screen. Cells were transduced with the lentiviral library at a low multiplicity of infection (0.3-0.4) by treating cells with 8 μg/mL polybrene and lentiviral supernatants for 24 hours. Transduced cells were selected by treatment with 20 μg/mL puromycin for 24 hours followed by trypsinization and reseeding in the same plates with 20 μg/mL puromycin for another 24 hours. After selection, cells were passaged for 6 days before splitting into untreated or OA, cisplatin, or Illudin S treated fractions. Cells were passaged for an additional 12 days with passaging every 3 days in medium with or without low doses of OA (1.7 nM), cisplatin (1 μM), and Illudin S (1.4 ng/mL) which corresponds to predetermined LD_20_ concentrations in uninfected cells. Cell pellets were harvested after completion of selection, which we consider the start of the screen, (T_0_) and at the final timepoint (T_18_).

#### Next generation sequencing and data analysis

Genomic DNA was extracted from treated and untreated cells at day 0 and day 18, and the genomic DNA regions containing the integrated sgRNAs were amplified by PCR using NEBNext Ultra II Q5 Master mix with the LCV2_forward and LCV2_reverse primers (Table S12). A second PCR reaction introduced i5 and i7 multiplexing barcodes (Table S12) and gel-purified PCR products were sequenced on Illumina NextSeq2000. Fastq files were generated using bcl2fastq v2.20.1, reads were trimmed to 20 bp using cutadapt 3.7 (58) and trimmed reads were assigned to guides in the TKOv3 library by MAGeCK 0.5.9.5 (59) to create a count matrix, where sequencing reads were mapped to the designed sgRNA library. To quantity relative sgRNA abundance, first, low-abundance sgRNAs (counts < 30) were excluded from downstream analysis to reduce noise and improve data reliability. Raw sequencing counts were then normalised to log2 transcripts per million (log2TPM) within each replicate to account for differences in sequencing depth. The log2TPM values were processed using the Linear Models for Microarray and Omics Data (limma) library (71) to define log2 fold changes in sgRNA abundance and subsequently converted to z-scores by comparing target sgRNAs against negative control (intergenic and non-targeting) sgRNAs. 4 key comparisons were performed: T_0_ vs. T_18_, to assess temporal changes in gRNA abundance over the course of the experiment; and untreated T_18_ vs. treatment (OA, cisplatin or Illudin S) at T_18_, to evaluate the behavior of guides in the three treatment conditions. A z-score cut-off of <-5 was set to identify gRNAs causing proliferation defects and synthetic lethality in OA, cisplatin or Illudin S. Log2 fold changes of gRNAs can be consulted through an interactive web portal at https://slim.icr.ac.uk/projects/base_editing_tiling.

#### Visualization of base editing data

*Linear residue maps* were generated in Python (3.12.2) using pandas (72), numpy 2.1.3 (73) and matplotlib 3.9.2 (74). Each plot shows the average log2 fold change of guides targeting a given residue. Domain and feature level annotations were gathered from UniProt and InterPro and mapped to the protein sequence.

#### Clustering of gene essentiality across treatments

was performed in Python (3.12.2) using pandas (72), numpy 2.1.3 (73) and matplotlib 3.9.2 (74). In summary, Pearson correlation coefficients were calculated for a pair of treatments for each gene, using log2 fold changes (treatment/untreated) of individual guides at T_18_. Treatment comparisons were okadaic acid vs. cisplatin, okadaic vs. illudin S, and cisplatin vs. illudin S. Only guides significant in at least one treatment condition (non-adjusted p-value < 0.05) were included. Using these, correlation matrices for all genes across treatments were calculated and used for hierarchical clustering of the genes using the Ward method (75) and Euclidean distance metric.

#### Interpretation and visualization of base editing data on protein structures and interaction site annotations

were performed in ChimeraX (76), using published crystal structures where available, or alternatively AlphaFold3 (AF3) prediction models (48). PP2A C subunit annotations were performed on PDB:4LAC; LCMT1 annotations were performed on PDB:3P71; B56epsilon annotations were performed on 8UWB with the KIF4A peptide modeled from PDB:6VRO; SPTLC2 annotations were performed on 7K0J. RIC8A GNA12 dimer models were generated in AlphaFold3 using protein sequences downloaded from Uniprot (RIC8A_HUMAN, Q9NPQ8; GNA12_HUMAN, Q03113-2), using GTP as ligand (ipTM = 0.82 pTM = 0.72). These predictions were overlayed with available crystal structures (RIC8A PDB:6TYL; GNA12 1ZCA) for model validation and interpretation. AlpaFold3 monomer (CYB5R4) and dimer models (CYB5R4-PP4 catalytic and CYB5R4-PP6 catalytic) were run on the AlphaFold server (https://golgi.sandbox.google.com/) (48), using protein sequences extracted form Uniprot, and co-factors and catalytic site metal ions available through the AlphaFold server. Specifically, protein sequences used were CYB5R4_HUMAN (<u>Q7L1T6</u>), PP4C_HUMAN (<u>P60510</u>) and PPP6_HUMAN (<u>O00743</u>). Co-factors used were heme (HEM), Flavin-adenine dinucleotide (FAD) Nicotinamide-adenine-dinucleotide-phosphate (NAP). Catalytic metal ions used were Fe and Zn. Seed number was set to auto. All models returned an interface predicted Template Modeling score (ipTM) and predicted Template Modeling score (pTM) above 0.5, which was used as cut-off for confidence (48,77,78) :

CYB5R4-HEM-FAD-NAP PP4-Fe Zn : iPTM = 0.65; PTM = 0.53

CYB5R4-HEM-FAD-NAP PP6-Fe Zn: iPTM = 0.68; PTM = 0.55

CYB5R4^1-153^-HEM: iPTM = 0.87; PTM = 0.74

CYB5R4^1-153^-HEM - PP4-Fe Zn: iPTM = 0.87; PTM = 0.88

CYB5R4^1-153^-HEM – PP6-Fe Zn: iPTM = 0.86; PTM =0.86

Best CYB5R4-phosphatase models were obtained with CYB5R4^1-153^-HEM (including the Cytb5 domain, omitting the CS and Cytb-R domains), displaying low predicted alignment error (PAE) scores (Sup Figure S5C), and high (i)pTMs up to 0.88. For each AF3 prediction, the 5 output models were aligned to assure consistency when interpreting base editing data in ChimeraX (76). Base editing data and interaction site annotations were performed using the AF3 model with highest ranking score (model_0).

### Validation of edits at CYB5R4 loci

Selected individual sgRNAs targeting CYB5R4 (H89R-targeting gRNA: TATCATCCTGGTGGAGAAGA; H112R-targeting gRNA: GTTCATCGTTGGGTCAATT; W114R-targting gRNA: ACCCAACGATGAACCTGGTA) were synthesized as DNA oligonucleotides containing Esp3I cloning overhangs: Forward oligo: 5’-CACCG[gRNA (20nt)]-3’; Reverse oligo: 5’-AAAC[reverse complement gRNA (20nt)]C-3’. A non-targeting sgRNA was used as control. The oligonucleotides were annealed and phosphorylated by T4 PNK (NEB) followed by golden gate cloning into pRDA_479 (40) with Esp3I and T7 ligase. The plasmids were transformed into NEB Stable Competent E. coli, which were grown on LB agar containing ampicillin at 30°C for 24 h. Plasmid DNA was prepared from single colonies, and gRNA insertion was validated by sequencing. Virus particles for each sgRNA expressing base editor construct were produced as described above. RPE1-hTERT p53-/-cells were transduced and selected as described above. After completion of selection, cells were cultured for 3 days (t3) before cell pellets were collected except for guide number 3, for which the cell pellet was harvested at t9 due to an insufficient number of cells at t3. Genomic DNA was extracted using the QIAamp DNA Blood Mini Kit (Qiagen). PCR reactions were set up to amplify the genomic CYB5R4 loci, where editing was predicted to occur, and introduce Illumina TruSeq Adaptor flaps. This was done using NEBNext Ultra II Q5 Master mix (New England Biolabs) with primers listed in Table S12. A second PCR reaction introduced Illumina i5 and i7 multiplexing barcodes using NEBNext Ultra II Q5 Master mix and Illumina TruSeq primers listed in Table S12. Gel-purified products were sequenced on Illumina NextSeq2000 and analyzed by CRISPResso2 (79).

### Yeast

#### Yeast strains & culture conditions

The *S. cerevisiae* strains used in this study were haploid and congenic to the S288C strain (BY4741). A list of yeast plasmids and yeast strains used in this study is provided in Table S13 and S15. Deletion of genes (*irc21* Δ*, rrd1* Δ and *sit4* Δ) and the C-terminal tagging of Sap185 was carried out by transformation of PCR-amplified cassettes as described previously (80) (list of primers Table S12). N-terminal tagging of Sit4 and Pph3 was achieved by the genomic integration of a *URA3* plasmid containing the native promoter and an N-terminally tagged version of the gene (plasmids have been constructed either by fusion PCR or ordered at GENESCRIPT and subcloned using restriction enzyme cloning; plasmid maps and plasmids are available upon request). Insertion of the construct at the respective endogenous locus was promoted by cutting the ORF in the plasmid with restriction enzymes. After insertion, selection on 5-Fluorooretic acid (FOA) for *ura3*Δ cells forced *URA3* containing plasmid DNA removal out of the genome. Candidate cells containing solely the tagged gene version were identified by PCR and western blot, and proper locus reconstitution was verified by DNA sequencing. This approach resulted in single N-terminally tagged genes expressed under control of their native promoters at their endogenous loci. In the case of *HA-PPH21* (YJV 905) an integrative *LEU2* plasmid was used instead, as an already available strain harbouring the *URA3* marker at the locus of *PPH21* was available. In this case, selection on FOA led to the loss of both auxotrophic markers and a reconstitution of the locus. Before harvest, all strains were grown into exponential phase on YPD medium (1% w/v yeast extract, 2% w/v bacto-peptone and 2% w/v glucose). For in vivo complementation, wildtype and mutant alleles of myc-Irc21 (purchased from GENESCRIPT) and myc-CYB5R4 (this study, Jakob Nilsson lab) were cloned into a centromeric *HIS3* plasmid both under the control of the native yeast promoter (plasmids are described in Table S13) and transformed into a *SAP185-HA irc21*Δ strain (YJV 1241). Before harvest cells were grown into exponential phase (2×10^7^ cells/ml) on glucose-containing synthetic dropout medium (SD, -HIS) for selection of plasmid markers.

#### Preparation of yeast pellets for PIB MS

1 liter of wildtype (wt) (BY4741) or *irc21*Δ (YJV 1206) cells was grown in YPD to 3×10^7^/ml, harvested by filtration (0.4 µM pore filters), washed once with PBS and snap frozen in liquid nitrogen. Frozen cell pellets were broken up using the SPEX 6770 cryogenic grinder (settings: 7 cycles; 2’ grinding at power level 14 and 3’ of cooling). The frozen cell powder contained about 40mg of total protein and was kept at −80°C until further analysis.

#### Immunoprecipitation and phosphatase activity assays with malachite green

25ml of yeast cultures expressing either HA-Pph21, myc-Pph3 or HA-Sit4 in the wildtype (YJV 905, YJV 1225, YJV 1275) or in the *irc21* Δ background (YJV 1198, YJV1228, YJV1277) as well as the BY4741 control strain was grown in YPD to a density of 2×10^7^cells/ml, harvested by centrifugation at 1000x g, snap frozen in liquid nitrogen and taken up in 600µl Lysis buffer (1%Triton, 50mM MES pH 6,5, 150mM NaCl, 1mM EDTA, 2mM DTT, cOmplete^TM^ protease inhibitor cocktail Roche). Cells were lysed with 400µl glass beads in a FastPrep^TM^-24 5G (MP Biomedicals) bead beating homogenizer (1 cycle 45’’ PL 6,5). For immunoprecipitation 500µl of the lysate was incubated for 1h at 4°C together with 20µl of Protein-A Sepharose beads CL-4B (Cytiva #17-0780-01) crosslinked either to HA-tag antibody 16B12 (mouse, Biolegend #901515, for HA-Pph21 and HA-Sit4 IP) or myc-tag antibody 4A6 (for myc-Pph3 IP). Immunoprecipitates (IPs) were washed 1x with 1ml Lysis buffer and 3x with 1ml Wash buffer (50mM MES pH 6,5, 150mM NaCl, 1mM EDTA, 2mM DTT); 1’ 400x g at 4°C. After the last wash, IPs were incubated with 200µl wash buffer containing 5mM ascorbate on ice for 20’, centrifugated again, and taken up in 400µl assay buffer (50mM Tris, 150mM NaCl) supplemented with 5mM ascorbate. 40µl of the beads/assay buffer suspension was mixed with 10µl of 4mM KRpTIRR phospho-peptide (dissolved in assay buffer) and incubated at 30°C shaking. The reaction was stopped at specific time points by the addition of 50µl malachite green mastermix (20µl of Malachite Green Phosphate Kit #MAK307 Sigma-Aldrich + 30µl assay buffer) 80µl of the final reaction (100µl) was transferred into a 96 well plate and the absorption of the malachite green molybdate phosphate complex was measured at 600 nm in a BioTeK synergy H1 micro plate reader. The absorption measured for the immunoprecipitated phosphatases, the “Total IP activity” was background corrected based on the absorption measured for the immunoprecipitate from the control strain lysate. To calculate specific activities (activity/immunoprecipitated HA-tagged phosphatase) of immunoprecipitates, 25% of the immunoprecipitated HA-tagged phosphatases was boiled in 1x protein loading dye (Laemmli buffer), analysed by western blot and the levels of immunoprecipitated HA-tagged phosphatases were quantified as described below. Datasets from three independent experiments were normalized to the arithmetic sum of each individual experiment (81), and means and standard deviations for each time point were calculated thereafter.

#### In vitro rescue with CYB5R4 wildtype and mutant

25ml of yeast cultures expressing HA-Sit4 in the wt (YJV 1225) or *irc21D* background (YJV 1228) as well as the BY4741 control strain were harvested by centrifugation at 1000x g, snap frozen in liquid nitrogen and taken up in 600µl lysis buffer II (1% Triton, 50mM Tris, 150mM NaCl, cOmplete^TM^ protease inhibitor cocktail Roche). Cells were lysed with 400µl glass beads in a FastPrep^TM^-24 5G (MP Biomedicals) bead beating homogenizer (1 cycle 45’’ PL 6,5). For immunoprecipitation 500µl of lysate was incubated for 1 h at 4°C together with 20µl of Protein-A Sepharose beads CL-4B (Cytiva #17-0780-01) crosslinked to HA-tag antibody 16B12 (mouse, Biolegend #901515). Sepharose Beads were washed 1x with 1ml lysis buffer and 3x with 1ml assay buffer (50mM Tris, 150mM NaCl) by centrifugation for 1’ 400x g at 4°C. After the last wash, beads were incubated with 200µl assay buffer containing 5mM ascorbate on ice for 20’. In parallel 0,13 ng of purified bacterially expressed mammalian CYB5R4 1-153 (wt or H89A/H112A mutant) was incubated for 15 ‘at RT in 60µl of CYB5R4 incubation buffer (50mM Tris, 150mM NaCl, 0,2% Triton X-100, 1mM ascorbate) giving a final concentration of ∼125 pM CYB5R4 in the incubation buffer (respectively ∼ 50 pM in the final assay). 20µl of each IP suspension was mixed with 20µl of the ascorbate-activated CYB5R4 (wildtype or mutant) or with mock buffer and incubated for 5’ at RT in 96 well plates. Phosphatase reaction was started by addition of 10µl of freshly prepared 2mM DiFMUP (in 50mM Tris, 150mM NaCl, 5% DMSO; Invitrogen #D6567, Lot # 2729822) and enzyme kinetics were determined by measuring the emission at 445 nm (358nm excitation) at 37°C in a BioTeK synergy H1 plate reader. For each time point, the DiFMUP emission of the phosphatase reaction, was background corrected based on the emission measured for the immunoprecipitate from the control strain lysate. To calculate specific activities (activity/immunoprecipitated HA-tagged phosphatase) of immunoprecipitates, 25% of the immunoprecipitated HA-tagged phosphatases was boiled in 1x protein loading dye (Laemmli buffer), analysed by western blot and the levels of immunoprecipitated HA-tagged phosphatases were quantified as described below. Datasets from three independent experiments were normalized to the arithmetic sum of each individual experiment (81), and means and standard deviations for each time point were calculated thereafter.

#### Western blot analysis and quantification

Whole cell protein lysates or IPs were boiled for 5min at 95°C in protein sample buffer (Laemmli buffer). Samples were separated by SDS-polyacrylamide gel electrophoresis (8% for SAP185-HA, 12% for Irc21 and 10% for all other proteins; Sigma Aldrich #01708, Tris/glycin buffered) and blotted on nitrocellulose membrane (GE Healthcare, 0.2μm). Membranes were stained with PonceauS, and blocked with 3% non-fat dry milk (NFDM) in PBS-Tween-20 (0.05%) for 1h at RT. The membranes were incubated with primary antibody (for antibodies & dilutions see Table S14) in 0.5% NFDM/PBS-Tween-20 o/n at 4°C. Incubation with secondary peroxidase conjugated antibody (1:10000 in 0.5% NFDM/PBS-Tween-20) was performed for 2 hours at RT, followed by incubation with western blotting detection reagents (GE Healthcare or Bio-Rad) as suggested by the manufacturer. Signal acquisition was performed using the ChemiDoc^TM^ (Bio-Rad) system. When western blot quantification was required, serial dilutions of the sample with the strongest signal were loaded, and linear or logarithmic regression of the resulting signals was used to calculate relative ratios between samples.

#### Figure models

Figure models were generated using Biorender (https://www.biorender.com/)

## Supporting information

Supplementary figures with legend

## Data availability

Base editing data is available through https://slim.icr.ac.uk/projects/base_editing_tiling Proteomics data is deposited in MassIVE (MSV000097024) and ProteomeXchange (PXD060411).

## Acknowledgements

We would like to thank Daniel Durocher for providing the RPE1 CYB5R4 knockout and the RPE1 p53 -/- Cas9 cell lines and the protein production and characterization unit at NNF CPR for helping with cloning and protein production. We would like to thank C.C. and M.S.J. from the Davies lab for assisting in operating the oxygen control chamber, and S. P. from the Zhu lab for technical assistance. We would like to thank the genomics facility at SUND UCPH for helping with NGS and analysis. Work at NNF CPR is supported by a grant from the Novo Nordisk Foundation (NNF14CC0001). Work in J.N.s laboratory is supported by grants from the Novo Nordisk Foundation (NNF23OC0082227 and NNF20OC0065098) and the Danish Cancer Society (R269-A15586 and R352-A20757). The work of the E.O. lab was funded by service and royalty fees from antibody licensing agreements and the monoclonal antibody service facility of the Medical University of Vienna/Max Perutz Labs. Work in the Kettenbach lab is supported by NIH R35GM119455. Work in the lab Zhu lab is supported by the University of Kansas Medical Center School of Health Professions, the University of Kansas Alzheimer’s Disease Center, and the National Institute on Aging (P30 AG072973, PI: Swerdlow). N.E.D. is funded by a Cancer Research UK Senior Cancer Research Fellowship (C68484/A28159).

## Author contributions

Conceptualization: B.M., S.M.A., A.N.K., E.O., J.N. Methodology: B.M., S.M.A., J.V., N.E.D., A.N.K., E.O., J.N. Software: N.E.D., M.H.M. Validation: B.M., S.M.A., J.V., N.E.D, A.N.K. Formal analysis: B.M., S.M.A., J.V., N.E.D., A.N.K. Investigation: B.M., S.M.A., J.V., R.C.R., G.V., B.L.B., M.H.M., E.C.G., C.B.C., M.B.W., D.H.G.,H.Z.,N.E.D., A.N.K., J.N. Resources: H.Z., A.N.K., E.O., J.N. Data Curation: B.M., S.M.A., M.H.M., N.E.D., A.N.K. Writing - Original Draft: B.M., S.M.A., J.N. Writing - Review & Editing: B.M., S.M.A., N.E.D., A.N.K., E.O., J.N. Visualization: B.M., S.M.A., M.H.M., N.E.D., J.N. Supervision: B.M., S.M.A., J.N. Project administration: B.M., S.M.A., J.V., H.Z., A.N.K., E.O., J.N. Funding acquisition: H.Z., N.E.D., A.N.K., E.O., J.N.

## References

1. Hunter T. Why nature chose phosphate to modify proteins. Philos Trans R Soc Lond B Biol Sci. 2012 Sep 19;367(1602):2513–6.

2. Depaoli-Roach AA, Park IK, Cerovsky V, Csortos C, Durbin SD, Kuntz MJ, et al. Serine/threonine protein phosphatases in the control of cell function. Adv Enzyme Regul. 1994 Jan;34:199–224.

3. Shi Y. Serine/threonine phosphatases: mechanism through structure. Cell. 2009 Oct 30;139(3):468–84.

4. Hwang J, Lee JA, Pallas DC. Leucine Carboxyl Methyltransferase 1 (LCMT-1) Methylates Protein Phosphatase 4 (PP4) and Protein Phosphatase 6 (PP6) and Differentially Regulates the Stable Formation of Different PP4 Holoenzymes. J Biol Chem. 2016 Sep 30;291(40):21008–19.

5. Lyons SP, Greiner EC, Cressey LE, Adamo ME, Kettenbach AN. Regulation of PP2A, PP4, and PP6 holoenzyme assembly by carboxyl-terminal methylation. Sci Rep. 2021 Nov 29;11(1):23031.

6. Meeusen B, Janssens V. Tumor suppressive protein phosphatases in human cancer: Emerging targets for therapeutic intervention and tumor stratification. Int J Biochem Cell Biol. 2018 Mar;96:98–134.

7. Ramos F, Villoria MT, Alonso-Rodríguez E, Clemente-Blanco A. Role of protein phosphatases PP1, PP2A, PP4 and Cdc14 in the DNA damage response. Cell Stress. 2019 Feb 21;3(3):70–85.

8. Verbinnen I, Vaneynde P, Reynhout S, Lenaerts L, Derua R, Houge G, et al. Protein Phosphatase 2A (PP2A) mutations in brain function, development, and neurologic disease. Biochem Soc Trans. 2021 Aug 27;49(4):1567–88.

9. Janssens V, Goris J. Protein phosphatase 2A: a highly regulated family of serine/threonine phosphatases implicated in cell growth and signalling. Biochem J. 2001 Feb 1;353(Pt 3):417–39.

10. Guo F, Stanevich V, Wlodarchak N, Sengupta R, Jiang L, Satyshur KA, et al. Structural basis of PP2A activation by PTPA, an ATP-dependent activation chaperone. Cell Res. 2014 Feb;24(2):190–203.

11. Sents W, Ivanova E, Lambrecht C, Haesen D, Janssens V. The biogenesis of active protein phosphatase 2A holoenzymes: a tightly regulated process creating phosphatase specificity. FEBS J. 2013 Jan;280(2):644–61.

12. Stanevich V, Jiang L, Satyshur KA, Li Y, Jeffrey PD, Li Z, et al. The structural basis for tight control of PP2A methylation and function by LCMT-1. Mol Cell. 2011 Feb 4;41(3):331–42.

13. Xing Y, Li Z, Chen Y, Stock JB, Jeffrey PD, Shi Y. Structural mechanism of demethylation and inactivation of protein phosphatase 2A. Cell. 2008 Apr 4;133(1):154–63.

14. Wu CG, Zheng A, Jiang L, Rowse M, Stanevich V, Chen H, et al. Methylation-regulated decommissioning of multimeric PP2A complexes. Nat Commun. 2017 Dec 22;8(1):2272.

15. Fellner T, Lackner DH, Hombauer H, Piribauer P, Mudrak I, Zaragoza K, et al. A novel and essential mechanism determining specificity and activity of protein phosphatase 2A (PP2A) in vivo. Genes Dev. 2003 Sep 1;17(17):2138–50.

16. Olivieri M, Durocher D. Genome-scale chemogenomic CRISPR screens in human cells using the TKOv3 library. STAR Protoc. 2021 Mar 19;2(1):100321.

17. Swingle MR, Honkanen RE. Inhibitors of Serine/Threonine Protein Phosphatases: Biochemical and Structural Studies Provide Insight for Further Development. Curr Med Chem. 2019 Jul 25;26(15):2634–60.

18. Favre B, Turowski P, Hemmings BA. Differential Inhibition and Posttranslational Modification of Protein Phosphatase 1 and 2A in MCF7 Cells Treated with Calyculin-A, Okadaic Acid, and Tautomycin. J Biol Chem. 1997 May;272(21):13856–63.

19. Olivieri M, Cho T, Álvarez-Quilón A, Li K, Schellenberg MJ, Zimmermann M, et al. A Genetic Map of the Response to DNA Damage in Human Cells. Cell. 2020 Jul 23;182(2):481–496.e21.

20. Dobrowsky RT, Kamibayashi C, Mumby MC, Hannun YA. Ceramide activates heterotrimeric protein phosphatase 2A. J Biol Chem. 1993 Jul 25;268(21):15523–30.

21. Zhu D, Kosik KS, Meigs TE, Yanamadala V, Denker BM. Gα12 Directly Interacts with PP2A. J Biol Chem. 2004 Dec;279(53):54983–6.

22. Yamaguchi Y, Katoh H, Mori K, Negishi M. Gα12 and Gα13 Interact with Ser/Thr Protein Phosphatase Type 5 and Stimulate Its Phosphatase Activity. Curr Biol. 2002 Aug;12(15):1353–8.

23. Zhu H, Qiu H, Yoon HW, Huang S, Bunn HF. Identification of a cytochrome b-type NAD(P)H oxidoreductase ubiquitously expressed in human cells. Proc Natl Acad Sci U S A. 1999 Dec 21;96(26):14742–7.

24. Deng B, Parthasarathy S, Wang W, Gibney BR, Battaile KP, Lovell S, et al. Study of the individual cytochrome b5 and cytochrome b5 reductase domains of Ncb5or reveals a unique heme pocket and a possible role of the CS domain. J Biol Chem. 2010 Sep 24;285(39):30181–91.

25. Benson DR, Deng B, Kashipathy MM, Lovell S, Battaile KP, Cooper A, et al. The N-terminal intrinsically disordered region of Ncb5or docks with the cytochrome b5 core to form a helical motif that is of ancient origin. Proteins. 2024 Apr;92(4):554–66.

26. Wang Y, Niu Y, Zhang Z, Gable K, Gupta SD, Somashekarappa N, et al. Structural insights into the regulation of human serine palmitoyltransferase complexes. Nat Struct Mol Biol. 2021 Mar;28(3):240–8.

27. Li S, Xie T, Liu P, Wang L, Gong X. Structural insights into the assembly and substrate selectivity of human SPT-ORMDL3 complex. Nat Struct Mol Biol. 2021 Mar;28(3):249–57.

28. Peters KA, Rogers SL. Drosophila Ric-8 interacts with the Gα12/13 subunit, Concertina, during activation of the Folded gastrulation pathway. Mol Biol Cell. 2013 Nov;24(21):3460–71.

29. Tall GG, Krumins AM, Gilman AG. Mammalian Ric-8A (synembryn) is a heterotrimeric Galpha protein guanine nucleotide exchange factor. J Biol Chem. 2003 Mar 7;278(10):8356–62.

30. Xie J, Zhu H, Larade K, Ladoux A, Seguritan A, Chu M, et al. Absence of a reductase, NCB5OR, causes insulin-deficient diabetes. Proc Natl Acad Sci. 2004 Jul 20;101(29):10750–5.

31. Wang W, Guo Y, Xu M, Huang HH, Novikova L, Larade K, et al. Development of diabetes in lean Ncb5or-null mice is associated with manifestations of endoplasmic reticulum and oxidative stress in beta cells. Biochim Biophys Acta. 2011 Nov;1812(11):1532–41.

32. Tsherniak A, Vazquez F, Montgomery PG, Weir BA, Kryukov G, Cowley GS, et al. Defining a Cancer Dependency Map. Cell. 2017 Jul 27;170(3):564–576.e16.

33. Lyons SP, Jenkins NP, Nasa I, Choy MS, Adamo ME, Page R, et al. A Quantitative Chemical Proteomic Strategy for Profiling Phosphoprotein Phosphatases from Yeast to Humans. Mol Cell Proteomics MCP. 2018 Dec;17(12):2448–61.

34. Mariano NC, Rusin SF, Nasa I, Kettenbach AN. Inducible Protein Degradation as a Strategy to Identify Phosphoprotein Phosphatase 6 Substrates in RAS-Mutant Colorectal Cancer Cells. Mol Cell Proteomics MCP. 2023 Aug;22(8):100614.

35. Guénolé A, Srivas R, Vreeken K, Wang ZZ, Wang S, Krogan NJ, et al. Dissection of DNA damage responses using multiconditional genetic interaction maps. Mol Cell. 2013 Jan 24;49(2):346–58.

36. Ferrari E, Bruhn C, Peretti M, Cassani C, Carotenuto WV, Elgendy M, et al. PP2A Controls Genome Integrity by Integrating Nutrient-Sensing and Metabolic Pathways with the DNA Damage Response. Mol Cell. 2017 Jul 20;67(2):266–281.e4.

37. Lue NZ, Liau BB. Base editor screens for in situ mutational scanning at scale. Mol Cell. 2023 Jul 6;83(13):2167–87.

38. Koblan LW, Doman JL, Wilson C, Levy JM, Tay T, Newby GA, et al. Improving cytidine and adenine base editors by expression optimization and ancestral reconstruction. Nat Biotechnol. 2018 Oct;36(9):843–6.

39. Walton RT, Christie KA, Whittaker MN, Kleinstiver BP. Unconstrained genome targeting with near-PAMless engineered CRISPR-Cas9 variants. Science. 2020 Apr 17;368(6488):290–6.

40. Sangree AK, Griffith AL, Szegletes ZM, Roy P, DeWeirdt PC, Hegde M, et al. Benchmarking of SpCas9 variants enables deeper base editor screens of BRCA1 and BCL2. Nat Commun. 2022 Mar 14;13(1):1318.

41. Hanna RE, Hegde M, Fagre CR, DeWeirdt PC, Sangree AK, Szegletes Z, et al. Massively parallel assessment of human variants with base editor screens. Cell. 2021 Feb 18;184(4):1064–1080.e20.

42. Richter MF, Zhao KT, Eton E, Lapinaite A, Newby GA, Thuronyi BW, et al. Phage-assisted evolution of an adenine base editor with improved Cas domain compatibility and activity. Nat Biotechnol. 2020 Jul 1;38(7):883–91.

43. McClelland LJ, Zhang K, Mou TC, Johnston J, Yates-Hansen C, Li S, et al. Structure of the G protein chaperone and guanine nucleotide exchange factor Ric-8A bound to Gαi1. Nat Commun. 2020 Feb 26;11(1):1077.

44. Kreutz B, Yau DM, Nance MR, Tanabe S, Tesmer JJG, Kozasa T. A New Approach to Producing Functional Gα Subunits Yields the Activated and Deactivated Structures of Gα_12/13_ Proteins. Biochemistry. 2006 Jan 1;45(1):167–74.

45. Freeman AK, Monteiro AN. Phosphatases in the cellular response to DNA damage. Cell Commun Signal CCS. 2010 Sep 22;8:27.

46. Nakada S, Chen GI, Gingras AC, Durocher D. PP4 is a gamma H2AX phosphatase required for recovery from the DNA damage checkpoint. EMBO Rep. 2008 Oct;9(10):1019–26.

47. Tsekitsidou E, Wong CJ, Ulengin-Talkish I, Barth AIM, Stearns T, Gingras AC, et al. Calcineurin associates with centrosomes and regulates cilia length maintenance. J Cell Sci. 2023 Apr 15;136(8):jcs260353.

48. Abramson J, Adler J, Dunger J, Evans R, Green T, Pritzel A, et al. Accurate structure prediction of biomolecular interactions with AlphaFold 3. Nature. 2024 Jun 13;630(8016):493–500.

49. Salvi F, Trebacz M, Kokot T, Hoermann B, Rios P, Barabas O, et al. Effects of stably incorporated iron on protein phosphatase-1 structure and activity. FEBS Lett. 2018 Dec;592(24):4028–38.

50. Xiao H, Jedrychowski MP, Schweppe DK, Huttlin EL, Yu Q, Heppner DE, et al. A Quantitative Tissue-Specific Landscape of Protein Redox Regulation during Aging. Cell. 2020 Mar 5;180(5):968–983.e24.

51. Welsh CL, Madan LK. Protein Tyrosine Phosphatase regulation by Reactive Oxygen Species. Adv Cancer Res. 2024;162:45–74.

52. Miki H, Funato Y. Regulation of intracellular signalling through cysteine oxidation by reactive oxygen species. J Biochem (Tokyo). 2012 Mar;151(3):255–61.

53. Tonks NK. Protein tyrosine phosphatases: from genes, to function, to disease. Nat Rev Mol Cell Biol. 2006 Nov;7(11):833–46.

54. Namgaladze D, Hofer HW, Ullrich V. Redox control of calcineurin by targeting the binuclear Fe(2+)-Zn(2+) center at the enzyme active site. J Biol Chem. 2002 Feb 22;277(8):5962–9.

55. Santos CXC, Hafstad AD, Beretta M, Zhang M, Molenaar C, Kopec J, et al. Targeted redox inhibition of protein phosphatase 1 by Nox4 regulates eIF2α-mediated stress signaling. EMBO J. 2016 Feb 1;35(3):319–34.

56. Nishito Y, Usui H, Shinzawa-Itoh K, Inoue R, Tanabe O, Nagase T, et al. Direct metal analyses of Mn2+-dependent and -independent protein phosphatase 2A from human erythrocytes detect zinc and iron only in the Mn2+-independent one. FEBS Lett. 1999 Mar 19;447(1):29–33.

57. Vit G, Duro J, Rajendraprasad G, Hertz EPT, Holland LKK, Weisser MB, et al. Chemogenetic profiling reveals PP2A-independent cytotoxicity of proposed PP2A activators iHAP1 and DT-061. EMBO J. 2022 Jul 18;41(14):e110611.

58. Martin M. Cutadapt removes adapter sequences from high-throughput sequencing reads. EMBnet.journal. 2011 May 2;17(1):10.

59. Li W, Xu H, Xiao T, Cong L, Love MI, Zhang F, et al. MAGeCK enables robust identification of essential genes from genome-scale CRISPR/Cas9 knockout screens. Genome Biol. 2014;15(12):554.

60. Colic M, Wang G, Zimmermann M, Mascall K, McLaughlin M, Bertolet L, et al. Identifying chemogenetic interactions from CRISPR screens with drugZ. Genome Med. 2019 Aug 22;11(1):52.

61. Hart T, Moffat J. BAGEL: a computational framework for identifying essential genes from pooled library screens. BMC Bioinformatics. 2016 Apr 16;17:164.

62. Ran FA, Hsu PD, Wright J, Agarwala V, Scott DA, Zhang F. Genome engineering using the CRISPR-Cas9 system. Nat Protoc. 2013 Nov;8(11):2281–308.

63. Hughes CS, Moggridge S, Müller T, Sorensen PH, Morin GB, Krijgsveld J. Single-pot, solid-phase-enhanced sample preparation for proteomics experiments. Nat Protoc. 2019 Jan;14(1):68– 85.

64. Eng JK, Jahan TA, Hoopmann MR. Comet: an open-source MS/MS sequence database search tool. Proteomics. 2013 Jan;13(1):22–4.

65. Elias JE, Gygi SP. Target-decoy search strategy for increased confidence in large-scale protein identifications by mass spectrometry. Nat Methods. 2007 Mar;4(3):207–14.

66. Valot B, Langella O, Nano E, Zivy M. MassChroQ: a versatile tool for mass spectrometry quantification. Proteomics. 2011 Sep;11(17):3572–7.

67. Schwanhäusser B, Busse D, Li N, Dittmar G, Schuchhardt J, Wolf J, et al. Global quantification of mammalian gene expression control. Nature. 2011 May 19;473(7347):337–42.

68. Tyanova S, Temu T, Sinitcyn P, Carlson A, Hein MY, Geiger T, et al. The Perseus computational platform for comprehensive analysis of (prote)omics data. Nat Methods. 2016 Sep;13(9):731–40.

69. Grassetti A V., Hards R, Gerber SA. Offline pentafluorophenyl (PFP)-RP prefractionation as an alternative to high-pH RP for comprehensive LC-MS/MS proteomics and phosphoproteomics. Anal Bioanal Chem. 2017;409(19):4615–25.

70. Papke CM, Smolen KA, Swingle MR, Cressey L, Heng RA, Toporsian M, et al. A disorder-related variant (E420K) of a PP2A-regulatory subunit (PPP2R5D) causes constitutively active AKT-mTOR signaling and uncoordinated cell growth. J Biol Chem. 2021;296:100313.

71. Ritchie ME, Phipson B, Wu D, Hu Y, Law CW, Shi W, et al. limma powers differential expression analyses for RNA-sequencing and microarray studies. Nucleic Acids Res. 2015 Apr 20;43(7):e47–e47.

72. The pandas development team. pandas-dev/pandas: Pandas [Internet]. Zenodo; 2024 [cited 2025 Feb 11]. Available from: https://zenodo.org/doi/10.5281/zenodo.3509134

73. Harris CR, Millman KJ, Van Der Walt SJ, Gommers R, Virtanen P, Cournapeau D, et al. Array programming with NumPy. Nature. 2020 Sep 17;585(7825):357–62.

74. Hunter JD. Matplotlib: A 2D Graphics Environment. Comput Sci Eng. 2007;9(3):90–5.

75. Ward JH. Hierarchical Grouping to Optimize an Objective Function. J Am Stat Assoc. 1963 Mar;58(301):236–44.

76. Pettersen EF, Goddard TD, Huang CC, Meng EC, Couch GS, Croll TI, et al. UCSF ChimeraX: Structure visualization for researchers, educators, and developers. Protein Sci Publ Protein Soc. 2021 Jan;30(1):70–82.

77. Zhang Y, Skolnick J. Scoring function for automated assessment of protein structure template quality. Proteins Struct Funct Bioinforma. 2004 Dec;57(4):702–10.

78. Xu J, Zhang Y. How significant is a protein structure similarity with TM-score = 0.5? Bioinformatics. 2010 Apr 1;26(7):889–95.

79. Clement K, Rees H, Canver MC, Gehrke JM, Farouni R, Hsu JY, et al. CRISPResso2 provides accurate and rapid genome editing sequence analysis. Nat Biotechnol. 2019 Mar;37(3):224–6.

80. Veis J, Klug H, Koranda M, Ammerer G. Activation of the G_2_ /M-Specific Gene *CLB2* Requires Multiple Cell Cycle Signals. Mol Cell Biol. 2007 Dec 1;27(23):8364–73.

81. Degasperi A, Birtwistle MR, Volinsky N, Rauch J, Kolch W, Kholodenko BN. Evaluating strategies to normalise biological replicates of Western blot data. PloS One. 2014;9(1):e87293.

