## Supplementary figures with legend for "A functional map of phosphoprotein phosphatase regulation identifies an evolutionary conserved reductase for the catalytic metal ions"

**Supplementary figure legends**

**Figure S1**

A) Western blot analysis of cell extracts from U2OS parental (PAR) cells, *CYB5R4* knockout (KO) cells, and *CYB5R4* KO cells stably complemented with CYB5R4-venus wildtype (WT) with the indicated antibodies. GFP antibody recognizes venus.

B) Validation of OA sensitivity for *CYB5R4* knockout cells. A colony formation assay was conducted in presence or absence of 2 nM OA with U2OS parental cells, *CYB5R4* knockout cells and *CYB5R4* knockout cells stably complemented with full length CYB5R4-venus WT. Representative images of colonies are shown.

C) Quantification of colony survival assay shown in B. The survival is calculated as the relative number of colonies in OA to untreated and represents three independent experiments. Error bares depict standard deviations and the shown *p*-values are based on one-way ANOVA analysis with Tukey's multiple comparisons test.

D) Volcano plot comparing the phosphatase components captured on phosphatase inhibitor beads from U2OS *CYB5R4* knockout (KO) and *CYB5R4* KO cells stably expressing CYB5R4-venus (WT rescue). The KO data is the same as in Figure 1E. PP2A components are indicated in green, PP4 in blue and PP6 in pink. 'C' indicates catalytic subunit. PIB-MS, phosphatase inhibitor beads and mass spectrometry (PIB-MS)

E) Heatmap showing depletion (blue) or enrichment (red) of PPP components from PIB-MS analysis of RPE parental cells and *CYB5R4* KO cells. Color scale indicates log2 median normalized intensities.

F) Heatmap showing depletion (blue) or enrichment (red) of PPP components from PIB-MS analysis of MEFs isolated from WT or *CYB5R4* KO mice. Color scale indicates log2 median normalized intensities.

G) Heatmap showing depletion (blue) or enrichment (red) of PPP components from PIB-MS analysis of U2OS parental and *CYB5R4* KO cells cultivated under standard tissue culture conditions (21% oxygen) and under physiological oxygen (6%). Color scale indicates log 2 median normalized intensities

H) Western blot analysis of pT35 MOB1 levels from cell extracts of U2OS parental cells, *CYB5R4* KO cells, and *CYB5R4* KO cells stably complemented with CYB5R4-venus WT or H89A/H112A mutant.

I) Volcano plot comparing the interactomes of 3xFLAG-PP4 C, immunopurified from U2OS parental or *CYB5R4* KO cells and analyzed by mass spectrometry.

J) Volcano plot comparing the interactomes of 3xFLAG-PP6 C, immunopurified from U2OS parental or *CYB5R4* KO cells and analyzed by mass spectrometry.

**Figure S2**

A) Volcano plot showing representation of PPP components from PIB-MS analysis of *S. cerevisiae* wildtype (*wt*) and *irc21* deletion strains. The catalytic subunits of PP2A-like phosphatases are indicated with their human orthologues in brackets.

B) Western blot analysis of immunoprecipitated HA-tagged Sap185 from *S. cerevisiae wt* and *irc21* and *rrd1* deletion strains with the indicated antibodies.

C) OA sensitivity of CYB5R4 domain mutants. A colony formation assay was conducted in presence or absence of 2 nM OA with U2OS parental cells, *CYB5R4* knockout cells and *CYB5R4* knockout cells stably complemented with venus-CYB5R4 full length, 1-153 (Cytb5 domain), 154-521 (CS + Cytb5R domains), or 272-521 (Cytb5R). Representative images of colonies are shown.

D) Quantification of the colony survival assay shown in C. The survival is calculated as the relative number of colonies in OA to untreated and represents three independent experiments. Error bares depict standard deviations, and the shown *p*-values are based on one-way ANOVA analysis with Tukey's multiple comparisons test.

E) Volcano plot comparing the interactomes of CYB5R4$^{1-153}$-venus WT to venus-tag alone, immunopurified from HeLa cells and analyzed by mass spectrometry. Components of PP2A (green), PP4 (blue), and PP6 (pink) as well as CYB5R4 (orange) are indicated.

F) Western blot analysis with the indicated antibodies of extracts from *S. cerevisiae wt* and *irc21* deletion strains complemented with myc-Irc21 wild-type or H158A/H182A (HH/AA) mutant; human myc-CYB5R4$^{1-153}$ WT or H89A/H112A (HH/AA) mutant; or full length venus-CYB5R4.

**Figure S3**

A) Residue map of cell fitness effects for each target. Each protein sequence is represented from left to right, and the color gradient represents the average log2 fold changes of gRNAs targeting the indicated residue at the screen end point ($T_{18}$) vs the starting point ($T_0$). Blue values specify that mutation of the target residue causes depletion (fitness defect) under unperturbed proliferation and indicates red enrichment. Grey specifies residues not targeted. Established PP2A-like holoenzyme components and regulators are shown in black and candidate regulators in grey. ABE, Adenine base editing. LFC, log2 fold change. OA, okadaic acid.

B) Log2 fold change ($T_{18}/T_0$) of guides comparing untreated versus okadaic acid treated conditions for the candidate regulators. U, untreated.

**Figure S4**

A) Residue map of PP2A catalytic subunit showing average residue log2 fold changes from the essential base editing tiling screen (untreated $T_{18}/T_0$). X-axis depicts the amino acid residue targeted for mutation. The y-axis shows the average log2 fold changes of guides targeting that residue. Bar color represent the log2 fold change with depletion shown in blue and enrichment in red. Grey specifies residues that are not targeted.

B) Average log2 fold changes of residues of PP2A catalytic subunit annotated on the crystal structure of PP2A catalytic subunit and catalytic metal ions (selected from PDB:4LAC). Functional domains and catalytic site are annotated, and strongly scoring residues are indicated. Depleted residues are shown in blue, enriched residues shown in red. Grey specifies residues that are not targeted.

C) Residue maps of B56epsilon regulatory subunit showing average residue log2 fold changes from the synthetic lethality base editing tiling screen (okadaic acid/untreated $T_{18}$).

D) Average log2 fold changes of residues of B56epsilon regulatory subunit annotated on the crystal structure of B56epsilon (selected from PDB:8UWB). Substrate binding pocket is indicated, with the KIF4A peptide (grey) overlayed (selected from PDB:6VRO).

E) Residue maps of SPTLC2 showing average residue log2 fold changes from the synthetic lethality base editing tiling screen (okadaic acid/untreated $T_{18}$).

F) Average log2 fold changes of residues of SPTLC2 annotated on the crystal structure of SPTLC1/2 + ssSPTa (PDB:7KOJ). SPTLC2 is showed as surface, SPTLC1 (white) and ssSPTa (grey) are shown as cartoon. The ssSPTSa binding pockets is indicated and the I130 residue highlighted in bold is mutated in disease.

G) Residue maps of GNA12 showing average residue log2 fold changes from the synthetic lethality base editing tiling screen (okadaic acid/untreated $T_{18}$).

H) Average log2 fold changes of residues of GNA12 and RIC8A annotated on a AlphaFold3 model of GNA12 and RIC8A. RIC8A is showed as surface and only the C-terminus of GNA12 is shown as cartoon.

**Figure S5**

A) Residue map of cell fitness in the presence of cisplatin for each target. Each protein sequence is represented from left to right, and the color gradient represents the average log2 fold changes of gRNAs targeting the indicated residue at the screen end point ($T_{18}$)) in treated conditions vs untreated conditions. Blue values specify that mutation of the target residue causes depletion (fitness defect) under cisplatin perturbed proliferation and red indicates enrichment. Grey specifies residues not targeted. Established PP2A-like holoenzyme components and regulators are shown in black and candidate regulators in grey. ABE, Adenine base editing. LFC, log2 fold change.

B) Residue map of cell fitness in the presence of Illudin S for each target. Each protein sequence is represented from left to right, and the color gradient represents the average log2 fold changes of gRNAs targeting the indicated residue at the screen end point ($T_{18}$) in treated conditions vs untreated conditions. Blue values specify that mutation of the target residue causes depletion (fitness defect) under Illudin S perturbed proliferation and red enrichment. Grey specifies residues not targeted. Established PP2A-like holoenzyme components and regulators are shown in black and candidate regulators in grey.

C) Validation of gRNA editing outcome for CYB5R4. Deep sequencing of endogenous CYB5R4 loci after transduction with single gRNAs targeting H89 or W114, followed by CRISPResso2 analysis, shows the frequency of mutated alleles. Reference: the genomic sequence. The gRNA and PAM are indicated as well as the amino acid translation. PAM, protospacer adjacent motif.

D) Western blot analysis of cell extracts from U2OS parental (PAR) cells, *CYB5R4* knockout (KO) cells, and *CYB5R4* knockout (KO) cells stably complemented with CYB5R4-venus wildtype (WT), H89R, H112R, and W114R with the indicated antibodies. GFP antibody recognizes venus.

E) OA sensitivity of CYB5R4 screen mutants. A colony formation assay was conducted in presence or absence of 2 nM OA with U2OS parental cells, *CYB5R4* knockout cells and *CYB5R4* knockout cells stably complemented with CYB5R4-venus WT, H89R, H112R, and W114R. Representative images of colonies are shown.

F) Cisplatin sensitivity of CYB5R4 screen mutants. A colony formation assay was conducted as in C but in presence and absence of 1 μM cisplatin. Representative images of colonies are shown.

G) Quantification of colony survival assay shown in F. The survival is calculated as the relative number of colonies in cisplatin to untreated and represents three independent experiments. Error bares depict standard deviations and the shown *p*-value are based on one-way ANOVA analysis with Tukey's multiple comparisons test.

**Figure S6**

A) Position Alignment Error plots of AlphaFold3 model of CYB5R4$^{1\text{-}153}$ with PP6 C (left) and PP4 C (right). Å, Ångstrom.

B) AlphaFold3 model of CYB5R4$^{1\text{-}153}$ with PP4 C with central residues indicated.

C) Left: surface view of PP4 C extracted from the CYB5R4$^{1-153}$ - PP4 C the AlphaFold3 model, showing contact sites of CYB5R4$^{1-153}$ (purple) and CYB5R4$^{1-153}$ W114 and heme (orange). Functional domains and catalytic site annotations are based on (3, 10-14). Right: average log2 fold changes of residues of PP4 C catalytic subunit annotated on PP4 C. Residues that score and are contacted by heme and CYB5R4$^{1-53}$ are indicated. LFC, log2 fold change.

D) Western blot analysis of immunoprecipitated HA-Sit4 from *S. cerevisiae* wildtype and *IRC21* deletion strains with the indicated antibodiesd. Rrd1 is the orthologue of PTPA.

**Figure S7**

A) DiFMUP dephosphorylation assay measuring the activity of purified PP4 holoenzyme in presence or absence of pre-reduced CYB5R4$^{1-153}$ WT or H89A/H112A. The data describes three independent experiments, and error bars represent standard deviations.

B-D) DiFMUP dephosphorylation assay measuring the activity of immunopurified HA-Sit4 from wildtype and *irc21* deletion strains (C) complemented with CYB5R4$^{1-153}$ WT (D) or CYB5R4$^{1-153}$ H89A/H112A (E). The data describes three independent experiments, and error bars represent standard deviations.

E) DiFMUP dephosphorylation assay measuring the activity of purified PP4 holoenzyme in presence or absence of the indicated reducing agents at 1 mM. The data describes three independent experiments, and error bars represent standard deviations.

F) Full time course of the DiFMUP dephosphorylation assay shown in Fig. 4F.

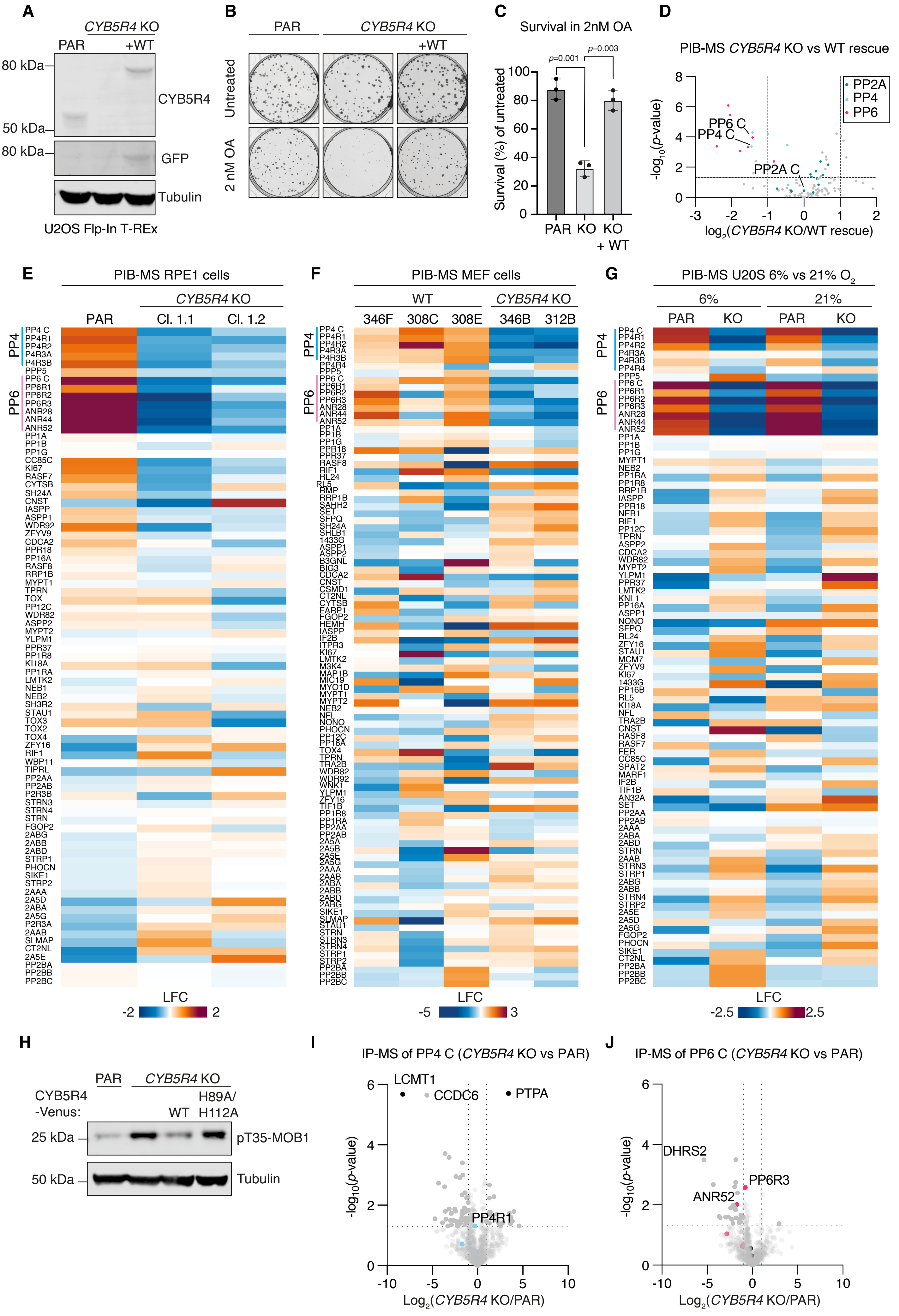


**Supplementary Figure 1**

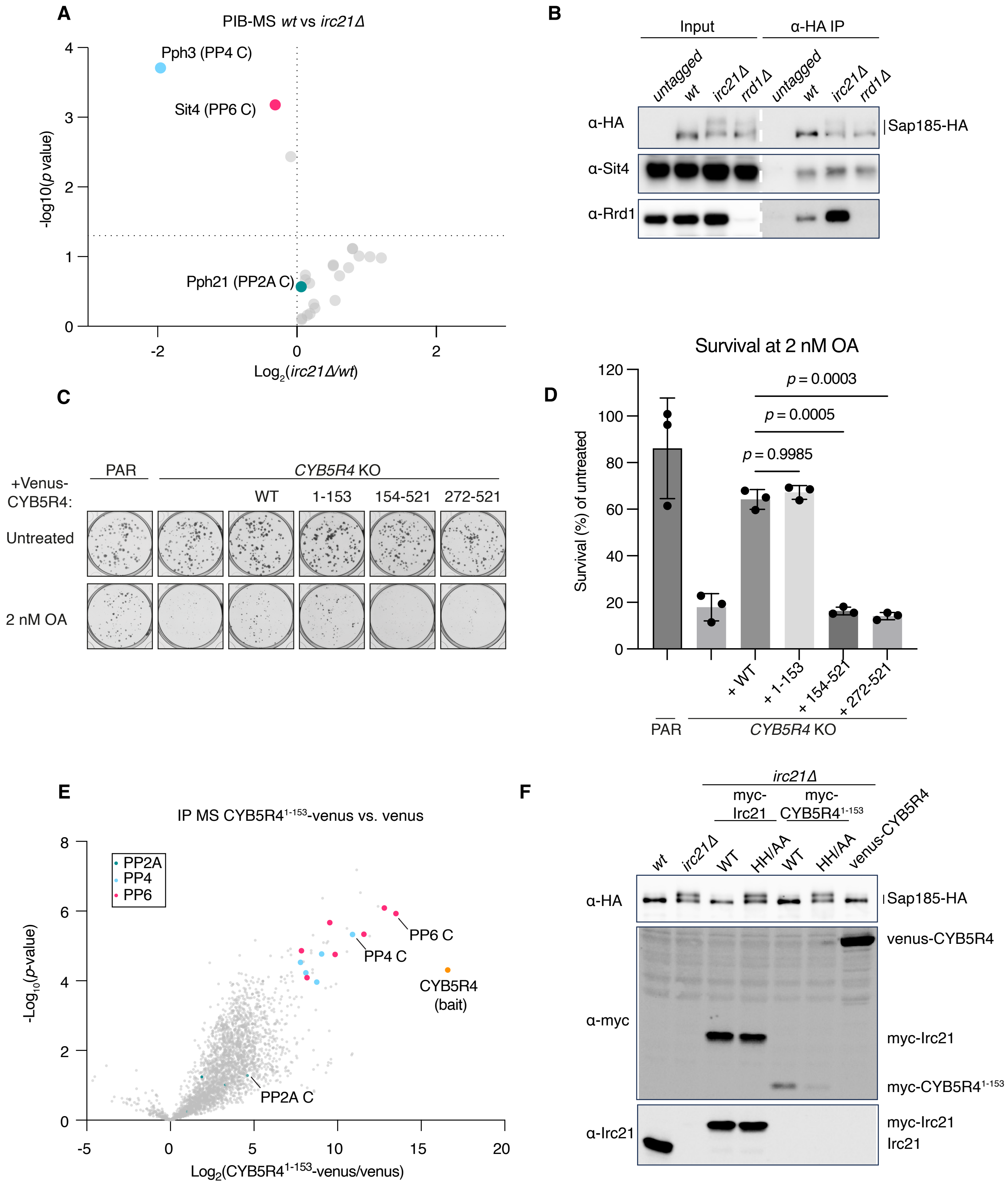


**Supplementary Figure 2**

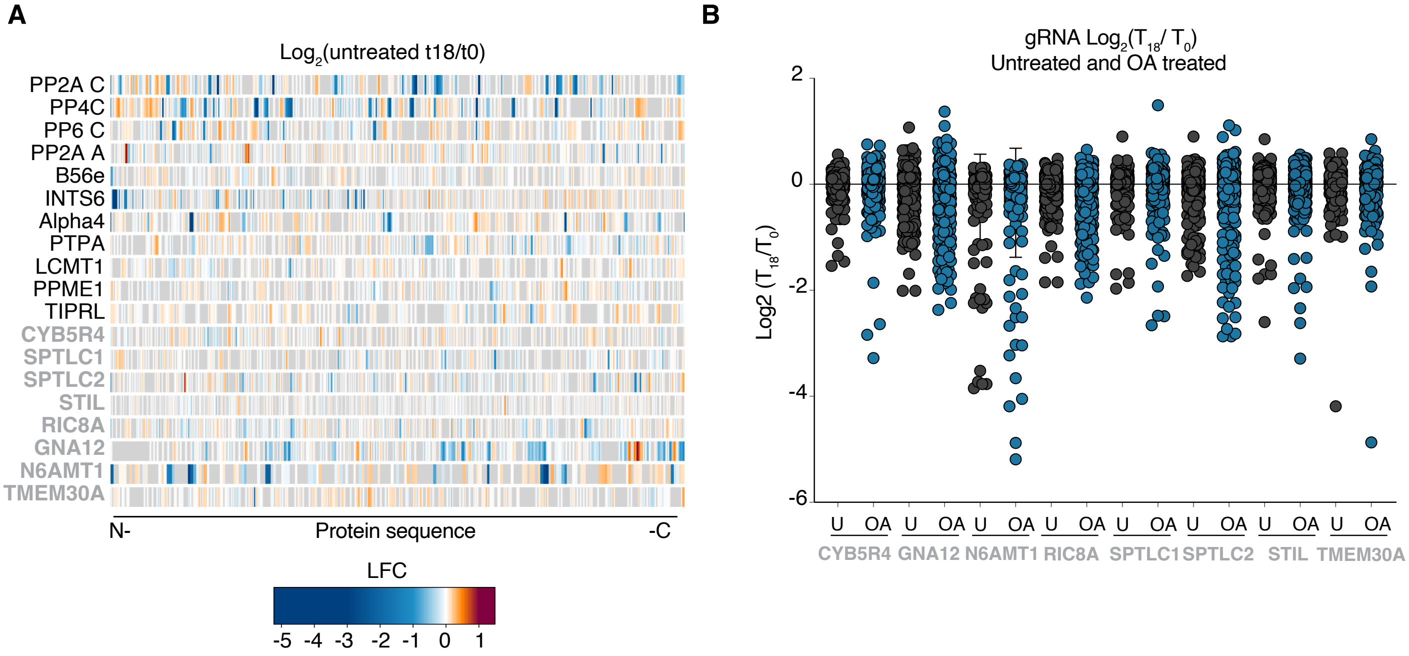


**Supplementary Figure 3**

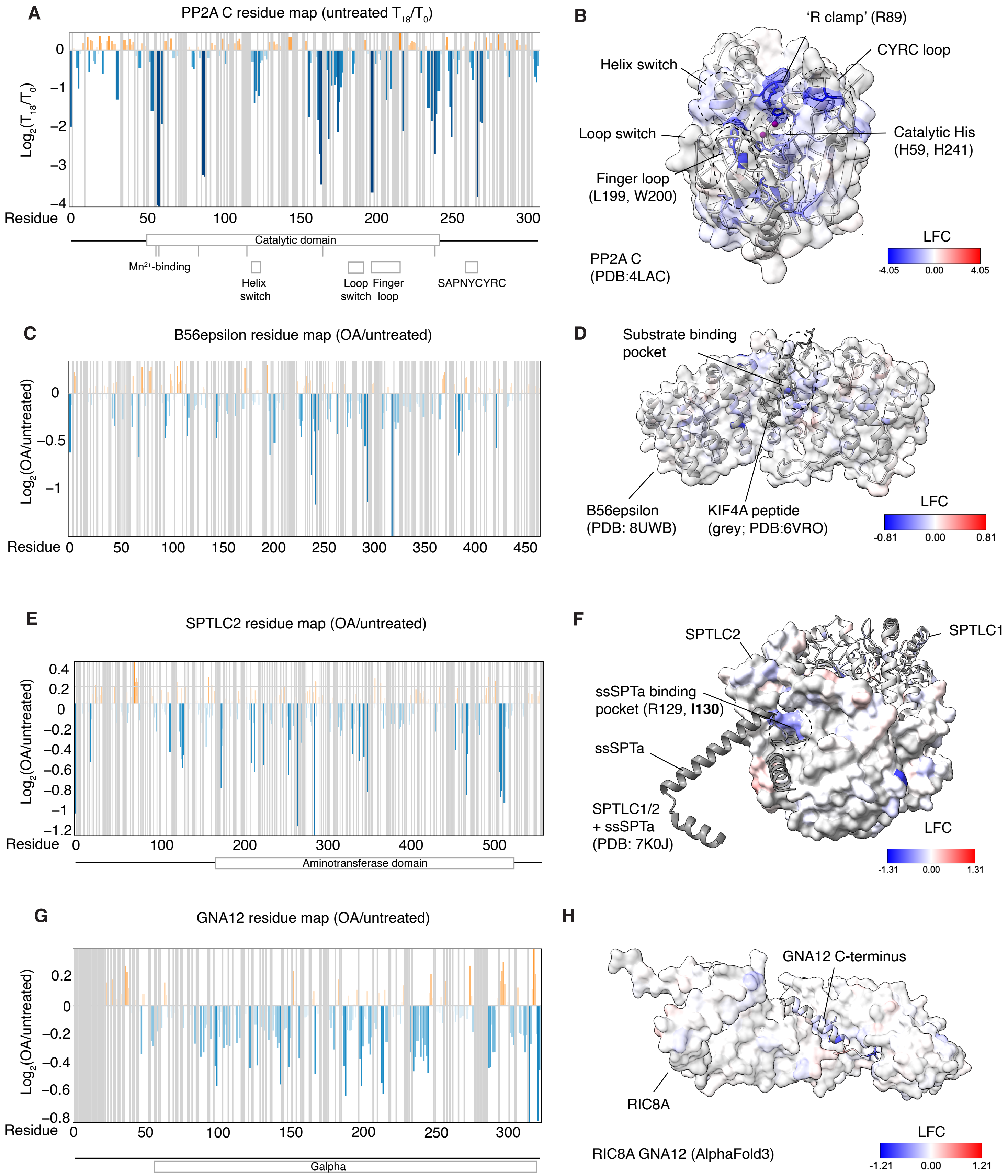


**Supplementary Figure 4**

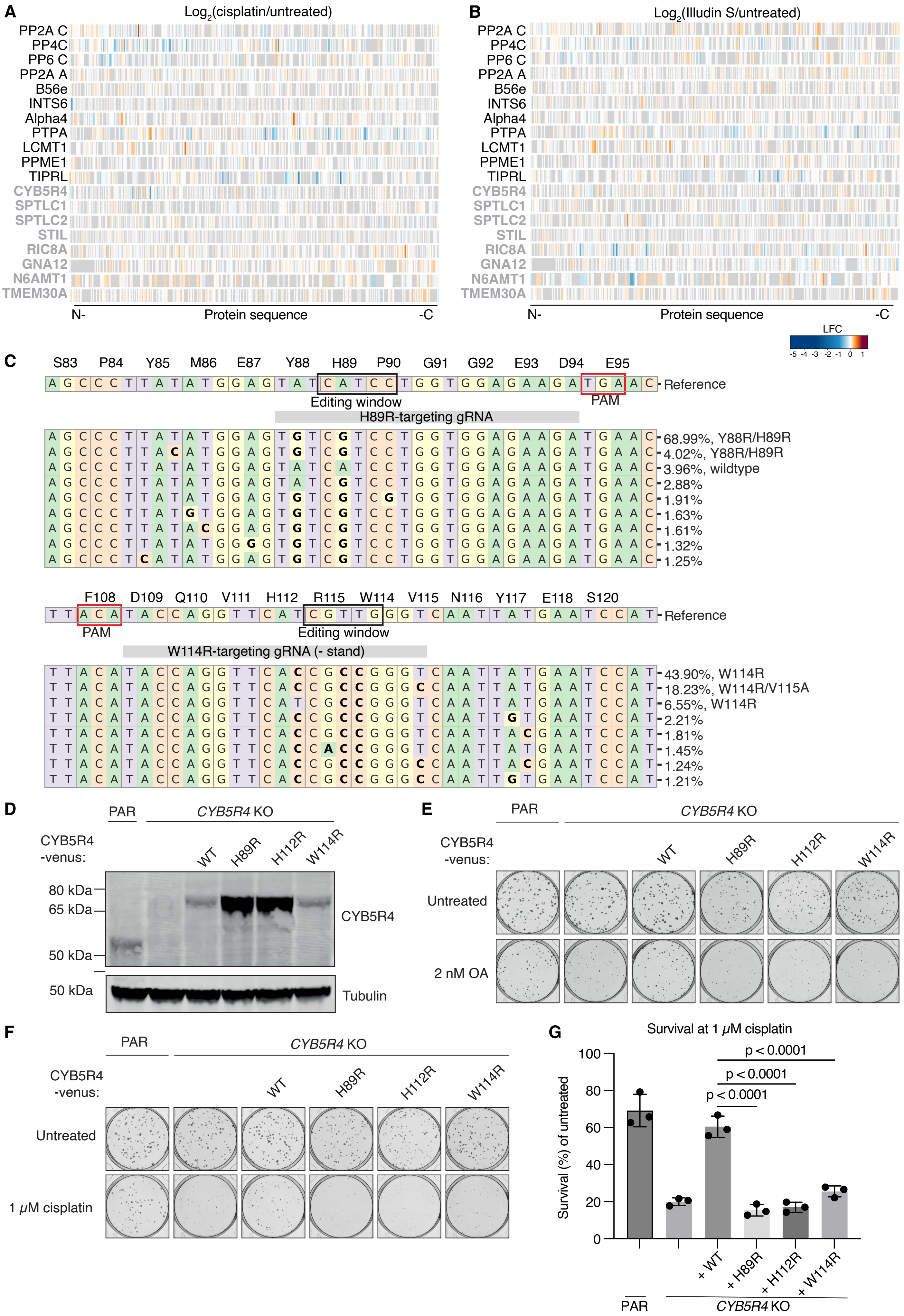


**Supplementary Figure 5**

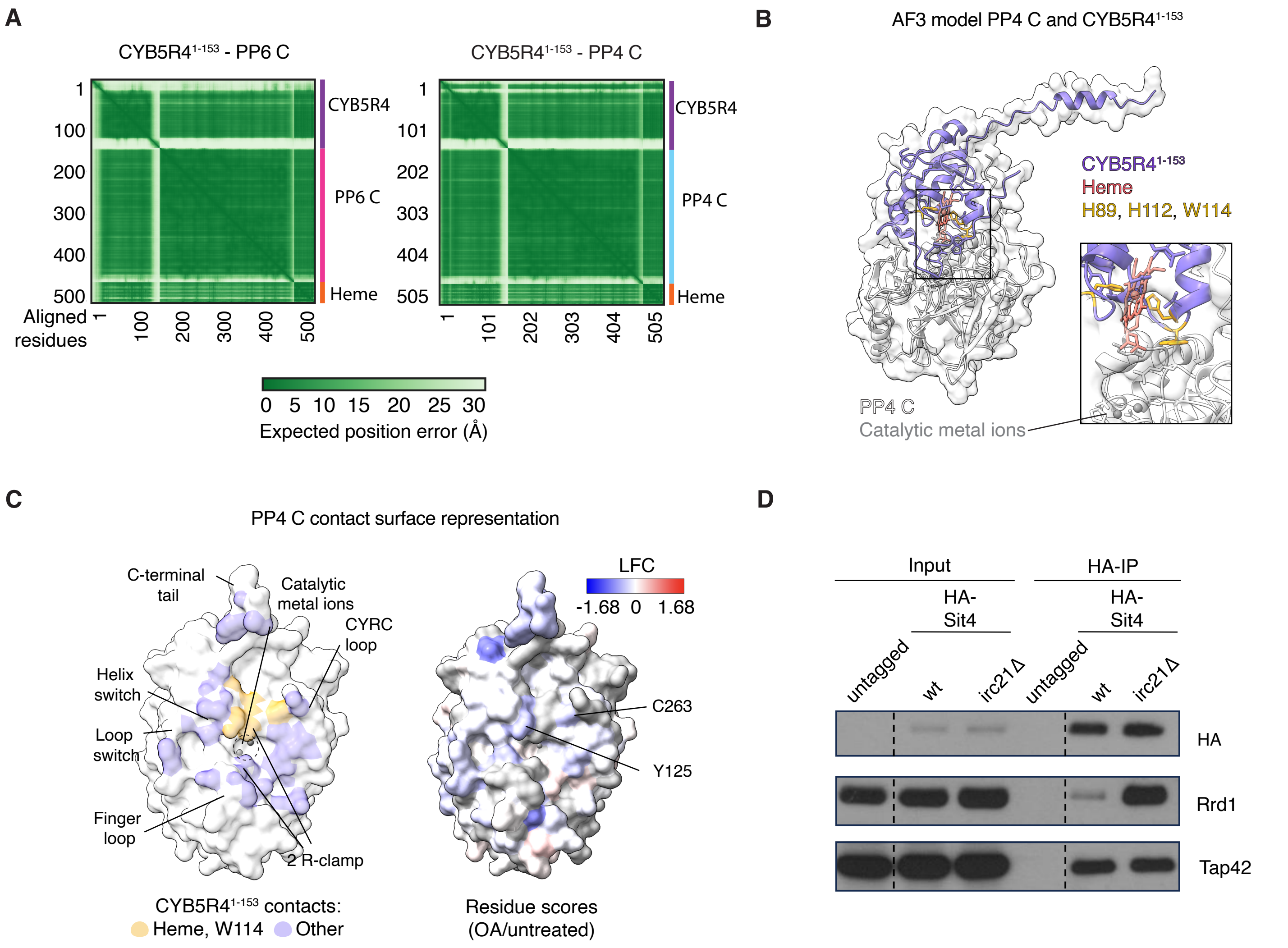


Supplementary Figure 6

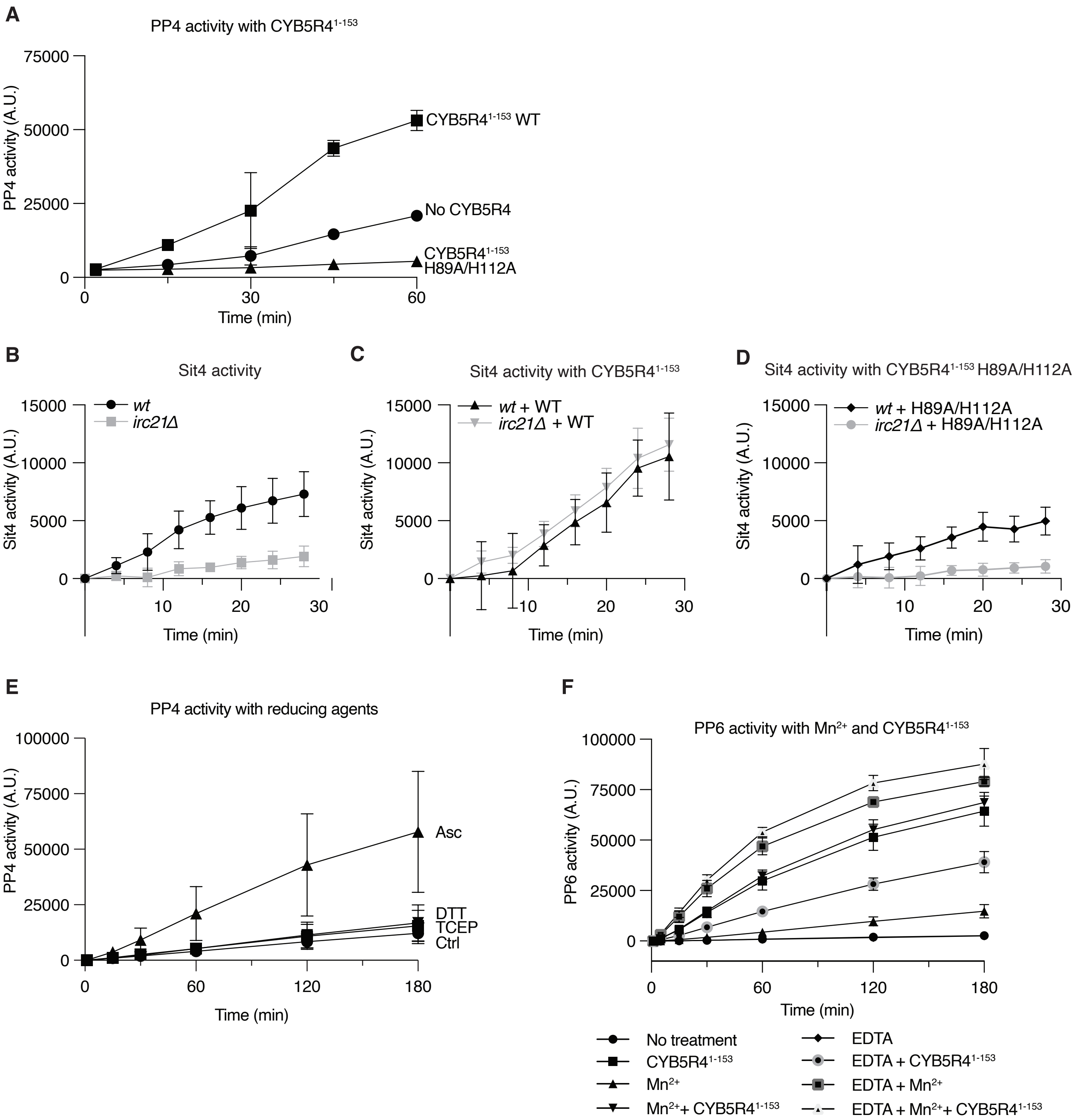


**Supplementary Figure 7**
